# Adversarial erasing enhanced multiple instance learning (siMILe): Discriminative identification of oligomeric protein structures in single molecule localization microscopy

**DOI:** 10.1101/2025.09.29.679377

**Authors:** Christian Hallgrimson, Y. Lydia Li, Claire A. Shou, Ben Cardoen, John Lim, Timothy Wong, Ismail M. Khater, Ivan Robert Nabi, Ghassan Hamarneh

**Affiliations:** School of Computing Science, Simon Fraser University, Burnaby, BC, Canada V5A 1S6; Department of Cellular & Physiological Sciences, Life Sciences Institute, University of British Columbia, Vancouver, BC, Canada V6T 1Z3; School of Biomedical Engineering, University of British Columbia, Vancouver, BC, Canada V6T 1Z3; Department of Electrical and Computer Engineering, Faculty of Engineering and Technology, Birzeit University, Birzeit, Palestine, P627

**Keywords:** single molecule localization microscopy, multiple instance learning, weakly supervised learning, caveolae, clathrin

## Abstract

Single-molecule localization microscopy (SMLM) achieves nanoscale imaging of complex protein structures in the cell. However, the ability to capture structural variability across cell conditions (cell lines, gene expression, treatment) from 3D point cloud SMLM data remains limited. We present siMILe, a weakly-supervised multiple instance learning (MIL) machine learning method to close this gap in interpretable subcellular discovery. siMILe identifies condition-specific changes in protein assemblies by leveraging their shape and network features, without requiring structure-level supervision. siMILe improves structure classification by extending embedded instance selection (MILES) through adversarial erasing and a symmetric classifier. We validated siMILe by detecting caveolae from caveolin-1 (Cav1) labeled PC3 prostate cancer cells differentially expressing cavin-1. In PC3-CAVIN1 cells, cavin-1 closely associates with siMILe-identified caveolae, to a lesser extent with higher-order non-caveolar Cav1 scaffolds, but not small Cav1 oligomers corresponding to 8S complexes, supporting a role for progressive cavin-1 interaction in 8S complex oligomerization. We also validated siMILe on simulated SMLM data and in detecting inhibitor-induced structural variations within clathrin-coated pit data. These results highlight siMILe’s potential to identify differential molecular structures in distinct cell conditions. siMILe extends the SuperResNET SMLM software platform with the ability to detect interpretable structural differences across conditions.

## 1 Introduction

Single-molecule localization microscopy (SMLM) is a super-resolution microscopy technique that achieves superior resolution based on detection of the stochastic blinking of isolated fluorophores [1]. SMLM achieves 20 nm lateral resolution, compared to the ∼250 nm resolution limit encountered by conventional microscopy [2], with more recent SMLM-based approaches such as MinFlux achieving 1-3 nm 3D resolution [1, 3]. The application of machine learning algorithms to 3D SMLM has increased the efficacy and accuracy of the reconstruction of super-resolution images from point emitter frames and for background fluorescence classification [4, 5]. However, approaches for quantitative analysis of 2D or 3D point-cloud data remain limited. Cluster analysis methods, including statistical, Bayesian, density-based, correlation-, tessellation-, image-, and machine-learning based approaches, have been applied to SMLM data [6, 7, 8]. The SuperResNET network analysis software platform leverages the power of graph-based construction and advanced machine learning techniques to process and interpret the complex point cloud data generated by SMLM [9]. Previously, we applied SuperResNET to caveolin-1 (Cav1) point clouds to classify caveolae and smaller non-caveolar oligomeric structures, called scaffolds [10] and detected caveolae and three classes of scaffolds as well as structural changes induced by point mutations to the Cav1 scaffolding domain [9, 11, 12]. More recently, SuperResNET was able to detect changes to clathrin-coated pits induced by small molecule inhibitors of clathrin endocytosis, which were not observed by transmission electron microscopy, and effectively interpret MinFlux clathrin pits and vesicles [13]. SuperResNET analysis of publicly available 2D SMLM data for nucleoporin Nup96 effectively segmented nuclear pores and Nup96 corners and distinguished 2 modules within the corners at 10.7±0.1 nm distance, thereby achieving molecular resolution [14]. Therefore, SuperResNET represents a highly sensitive approach to studying the spatial or molecular architecture of subcellular structures in situ in the intact cell.

The primary strengths of SuperResNET lie in its ability to analyze the network structure inherent in SMLM data and its integration of advanced machine learning algorithms for pattern recognition and structure identification. These features enable SuperResNET to uncover subtle structural patterns and relationships that might be overlooked by conventional analysis methods. The use of prior knowledge, in the form of defined group labels, enabled identification by SuperResNET of diverse Cav1 structures showing that the base Cav1 oligomer (the so-called 8S complex) combines to form higher-order scaffolds and caveolae [9, 15]. PC3 prostate cancer cells that express Cav1, but not the adaptor protein cavin-1 (also called PTRF) required for caveolae formation, were compared with PC3-CAVIN1 cells transfected with cavin-1 that now present caveolae, defining parameters that enabled identification of known structures, the 8S complex and caveolae, and thereby previously unknown intermediate higher order oligomers [9]. SuperResNET has also been used to study the spatial relationship on caveolae of the adaptor proteins EHD2 and PASCIN2 [16].

Here we build on SuperResNET to present a discovery method named siMILe, a weakly-supervised, machine learning algorithm designed to identify discriminatory changes in mesoscale domain structures between conditions. The idea is to leverage only weak labels: the image- or cell-level information, such as cell type, epigenetic, or environmental interventions [17]. ‘Strong’ labels in this context would be object- or structure-level annotations, which in SMLM often are absent.

Such a problem statement is frequently tackled by multiple instance learning (MIL) methods. Unlike traditional supervised learning techniques, where labels are assigned to each object, or ‘instance’, with MIL, the assumption is that only a set, or collection of instances, called ‘bags’, can be given a label. This is particularly effective when individual instances cannot be labeled, whether infeasible or expensive. It is important to note that this differs from quantifying the structural diversity or heterogeneity [18, 19], given that discovery of diversity does not in itself leverage weak labels to find discriminative differences. MIL has been applied to many fields, ranging from confocal microscopy [20], drug efficacy discovery [21], DNA protein identification, and histopathology classification [13]. A more recent proposed MIL approach [22] optimizes its output so that nearby objects have the same or a similar object label. While this is a valid constraint in several MIL applications, here it would not be appropriate because in SMLM proximate protein complexes can be quite dissimilar.

In this paper, not only are we the first to apply MIL to SMLM, but we enhance MIL via embedded instance selection (MILES) [14] for improved instance classification through the first use of adversarial erasing in MIL, enhanced with a new symmetric classifier. Adversarial erasing iteratively removes identified structures and retrains the model, ensuring detection of all discriminative structures rather than just the most prominent ones. The symmetric classifier enables simultaneous identification of structures unique to each condition in a single analysis, eliminating the need for separate comparisons and improving computational efficiency. siMILe, illustrated in Fig 1, uses only image-level labels, allowing the potential for discovery by eliminating the need for manual annotations. The method is applied to a simulated dSTORM dataset to evaluate its efficacy, and to PC3 and PC3-CAVIN1 cells expressing cavin-1 to display the ability of siMILe to extract discriminative structures from SMLM. The trained model from this dataset is then applied to a new dataset of PC3-CAVIN1 cells dually labeled for Cav1 and cavin-1 to investigate its generalizability, and to verify using object level ground truth that the structures detected by siMILe are indeed discriminate from a biological perspective. This is achieved by validating the findings by contrasting them to protein interactions, information that was withheld from siMILe. We further extended the application of siMILe to clathrin-mediated endocytosis, identifying structural changes in clathrin-coated pits in response to small molecule inhibitors. These results support the ability of siMILe to identify changes in mesoscale domain structure, advancing our understanding of differential expression of intracellular molecular structures in distinct cell conditions.

**Figure 1.**
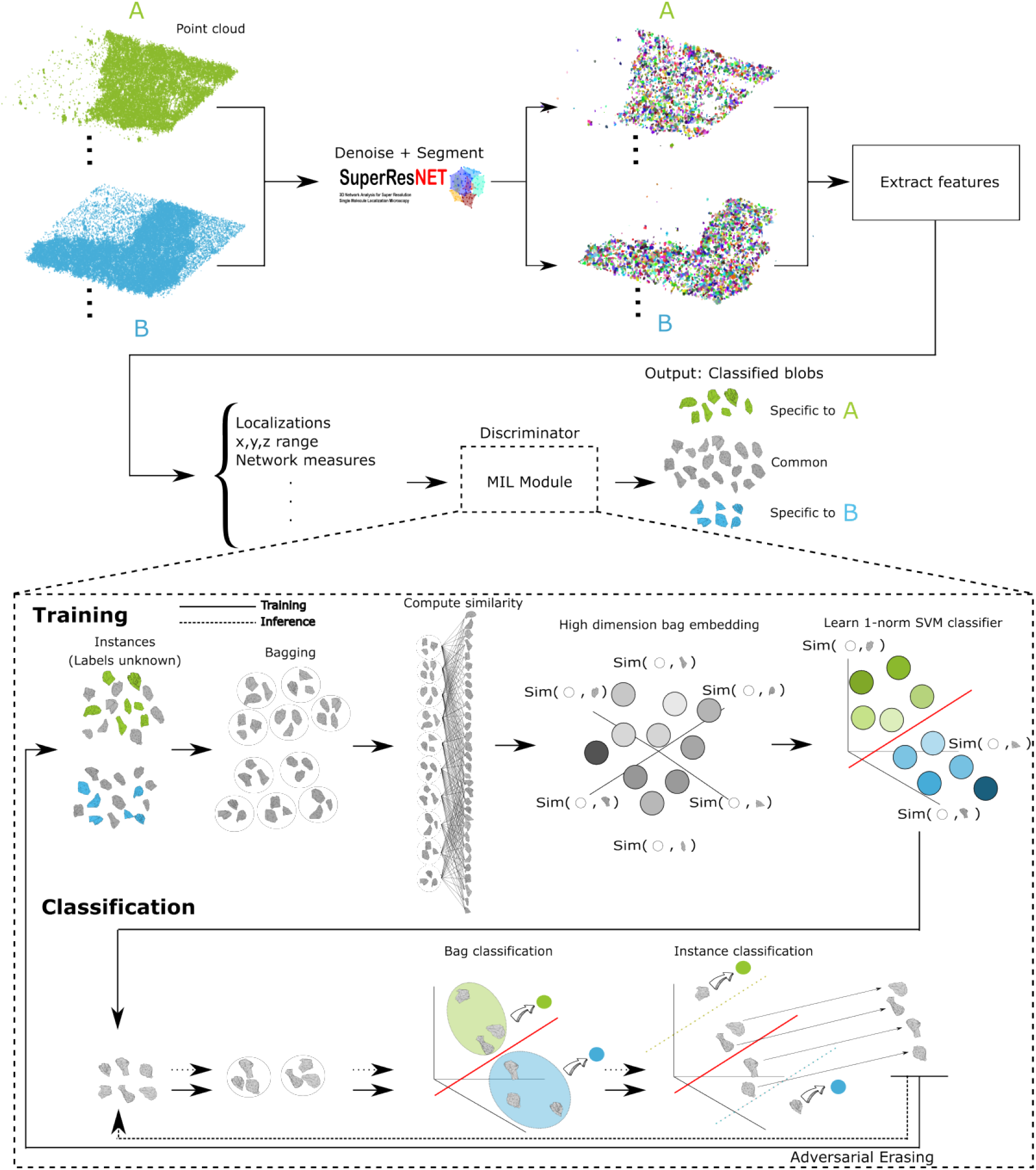
Overview of the proposed pipeline. The data preprocessing is shown at the top of and *B* into SuperResNET for denoising and segmentation, followed by feature extraction. Next, the processed data of both conditions are input into the siMILe module for classification into either structures specific to *A*, structures specific to *B*, or ambiguous. The details of siMILe are given within the dotted box. To train, the instances from a given class are grouped into bags, these bags are given an embedding by comparing all of its instances to the training data. Each dimension is represented by comparing the similarity of the bag to a given training instance. Next, a l1norm SVM classifier learns to fit a hyperplane that best separates the bags based on class. The instances of a bag can be labeled based on their contribution to the bags classification. Unlabeled instances are reused in the adversarial erasing steps, where the model is retrained using them. Structures are visualized using the convex hull of its localizations for claritythe figure, which includes inputting SMLM conditions *A*.

## 2. Results

### 2.1 Problem statement

In SMLM, each acquired image produces a 3D point cloud of molecular localizations. Throughout this paper, we use “image” and “point cloud” interchangeably to refer to these SMLM datasets. Formally, we are given input *I*: a set of *n* pairs (*P*_*i*_, *Y*_*i*_) for *i* = 1, …, *n*, where *P*_*i*_ is a point cloud and *Y*_*i*_ is an associated class label. Each point cloud *P*_*i*_ consists of (*x, y, z*) coordinates (3D vectors of real numbers), and the class labels *Y*_*i*_ ∈ {0, 1} indicate the condition from which the point cloud is derived.

Our expected output is a set of *n* pairs (*Q*_*i*_, *Ŷ*_*i*_) for *i* = 1, …, *n*. Each *Q*_*i*_ is a set of *J*_*i*_ disjoint subsets {*Q*_*ij*_ : *j* = 1, …, *J*_*i*_}, where each subset *Q*_*ij*_ ⊂ *P*_*i*_ represents a segmented structure and *Q*_*ij*_ ∩ *Q*_*ik*_ = ∅ for all *j* ≠ *k*. Each *Ŷ*_*i*_ is a set of labels {*Ŷ*_*ij*_ : *j* = 1, …, *J*_*i*_} corresponding to the subsets, where *Ŷ*_*ij*_ ∈ {0, 1, −1} indicates whether subset *Q*_*ij*_ is discriminative to condition 0, discriminative to condition 1, or common to both conditions (-1). Algorithm 1 summarizes the mathematical notation of the problem setup.

#### Algorithm 1 Problem formulation

for learning discriminative segmentations from point clouds with weakly supervised labels. The input is a set of 3D point clouds with a binary label for each point cloud based on its class. The output corresponding to a given input is a set of disjoint segmentations on the input, where each segmentation can be labeled as discriminative to the class given by the input label, or not discriminative (common).

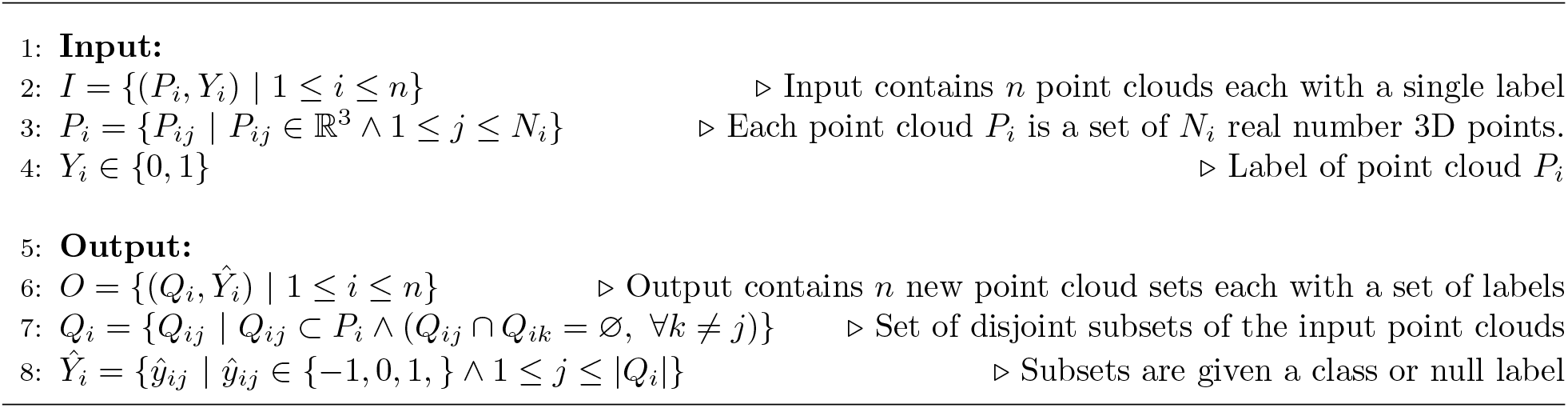

We use the SuperResNET network analysis platform to segment the point cloud, motivated by previous work supporting its ability to segment and extract clusters representing oligomeric biological structures [9, 11, 12]. A mean-shift algorithm segments the localizations into clusters, constellations of 3D points, or ‘blobs’ that represent these structures. SuperResNET also extracts features from these blobs based on size, shape, topology, point statistics, and graph networks. We refer the interested reader for more detail to the original work [9]. After SuperResNET processes all point clouds from a given condition, it produces segmented structures called ‘blobs’ (the *Q*_*ij*_ subsets). Each blob is represented by a 30-dimensional feature vector, yielding a feature matrix of shape *N* × 30 where *N* is the total number of blobs across all point clouds in that condition. We use this representation to compare conditions and extract blobs deemed distinct to their condition and are therefore discriminative. Next, we introduce multiple instance learning (MIL) as the core underlying weakly-supervised formulation used to learn the discriminative blobs in the output of our problem setup. We then introduce the diverse density (DD) MIL framework [23] that uses a single concept vector to classify bags of instances in feature space based on proximity to the concept vector. Next, we present multiple-instance learning via embedded instance selection (MILES) [24] which extends DD with the use of multiple concept vectors. This enables learning more complex relationships for classifying bag labels, and provides an algorithm for blob classification. Finally, we detail our method, siMILe, which extends MILES. The contributions of siMILe focus on the integration of adversarial erasing (AE) for improved instance classification and a symmetric classifier (SC) to reduce unnecessary computation time by asymmetric AE iterations. The base functionality of MIL algorithms is to identify discriminative from common objects, given two labels. But this can also cover the simpler use case where one set (A) or label is enclosed by the other one (B). In this case, there are no objects unique to A, only common to both and discriminative to B. In our problem statement, this case is not warranted; we need to be able to find those objects that are discriminative to each label individually. The combination of AE+SC in siMILe is designed to improve instance classification when discriminative instances exist in both classes (Fig. 1).

### 2.2 Multiple Instance Learning for SMLM

The problem formulation outlined in Algorithm 1 describes a weakly supervised learning paradigm where we need to extract discriminative blobs within an image using only the information provided by its condition label. Our method makes use of multiple instance learning (MIL), a weak supervision framework introduced by Dietterich et al. for drug activity prediction [25]. MIL identifies objects or ‘instances’ under conditions where class labels are represented by sets of instances, as opposed to individual instances. In the context of this paper, these ‘instances’ are the blobs (segmented structures) produced by SuperResNET, representing approximate protein structures. Training data points (the instances/blobs in our application) are grouped into bags containing a set number of instances. Using these bags, where each bag has a label that corresponds to a condition label under which the image is acquired, the focus of some MIL methods (e.g., MI-SVM [26], Citation-kNN [27], EM-DD [28]) is predicting a bag label using the aggregated information of its instances.For other methods (e.g., mi-SVM [26], APR [25]), it is more important to classify the individual instances by taking advantage of the representations learned by predicting their bag. Aggregating the information of the instances is useful when some instances are not capable of representing their weak label, while others are, and this distinction is unknown. In the traditional MIL formulation, there are two bag classes: positive and negative. The positive class contains instances that can be labeled as positive or negative, whereas the negative class is assumed to only contain negative instances. This 2-class setup is visualized in Figure 2-A. Given that an instance of the positive class has the potential to be found in the negative class, classification is performed on bags. In positive bags, the proportion of positive instances it contains is referred to as the witness rate [29]. In the traditional formulation, it is assumed that there is at least one positive instance in the bag; otherwise, a positive bag would be indistinguishable from a negative bag.

**Figure 2.**
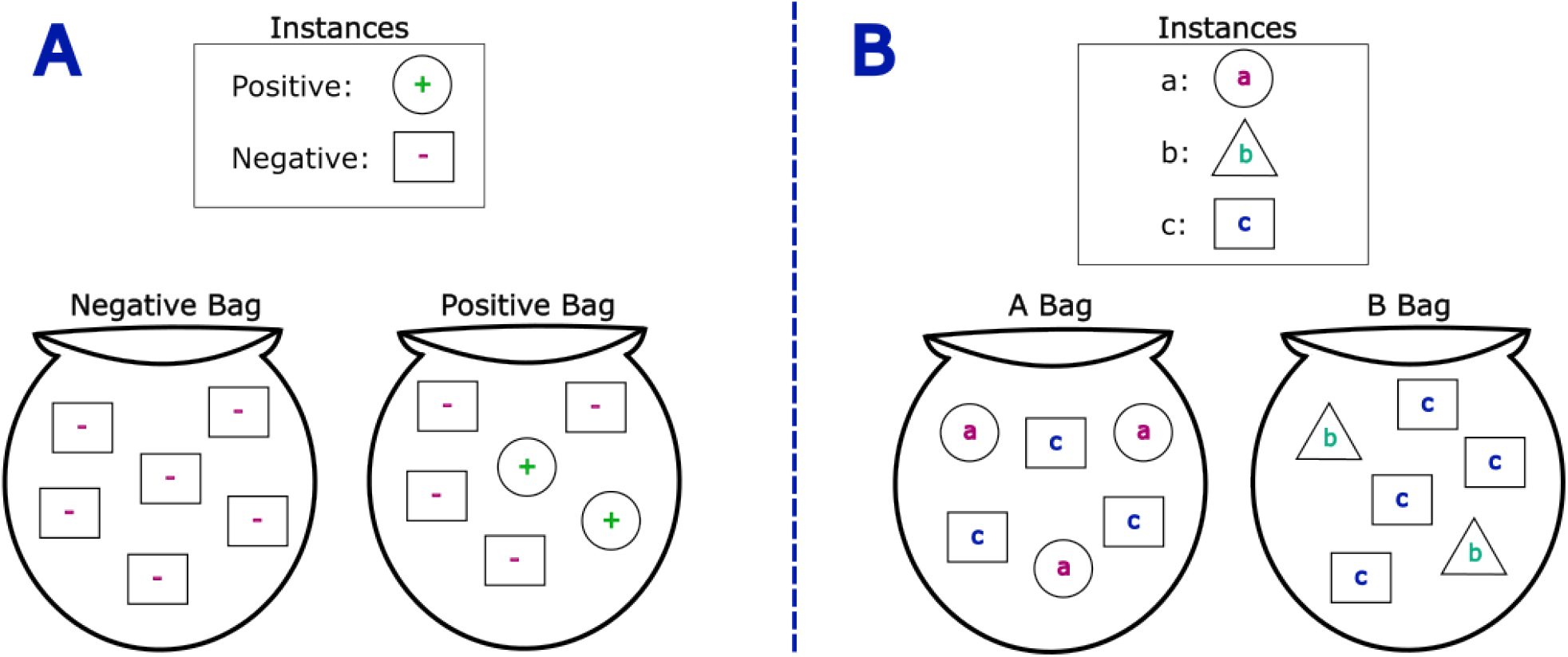
Visualizing two different multiple instance learning formulations. (A) Depicts the traditional MIL formulation with two classes: positive and negative. Both classes consist of negative instances, while only the positive class contains positive instances. Therefore, it is assumed that while a positive bag will have at least one positive instance, a negative bag will have only negative instances. (B) The case where both classes contain instances not found in the other. Class *A* contains instances of type *a* and *c*, while class *B* contains instances of type *b* and *c*. Thus, it is assumed that while each class contains instances shared with the other, they also contain instances that are specific to their class.

### 2.3 MILES

Diverse density (DD) [30] is a MIL algorithm notable for the use of a single ‘concept’ to aid in classifying bags, where this concept is a *representation of the relationship between positive and negative classes*. The target concept is assumed to be closer to positive instances and further away from negative ones, providing a method to differentiate the bags according to their label, as seen in Figure 3. Given a potential concept *c*, positive bag 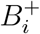, and negative bag 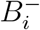, the concept that is most likely to fit the bags is determined by the number of positive and negative bags that agree on the concept by maximizing:

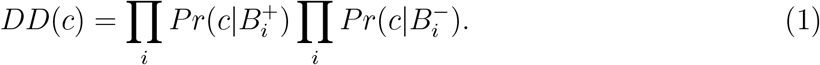

**Figure 3.**
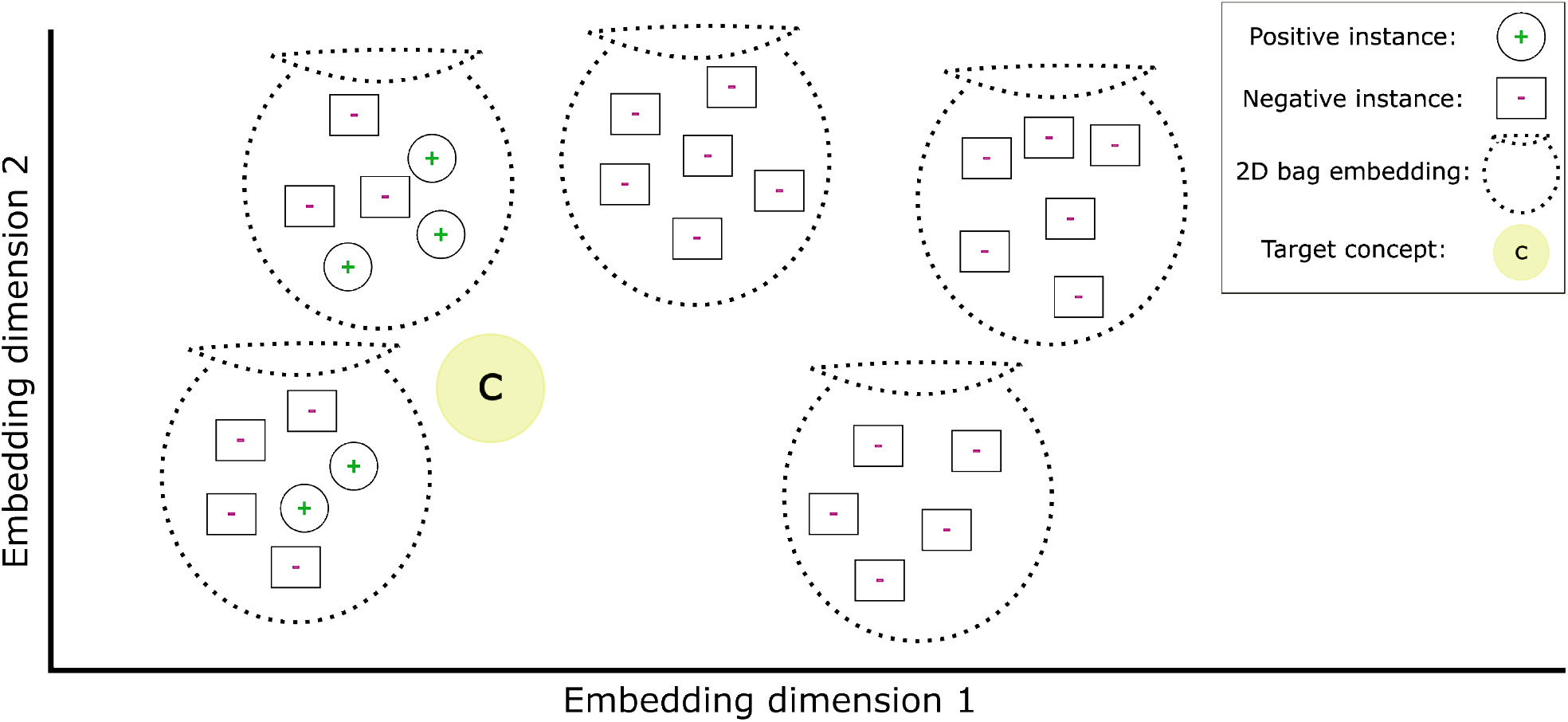
Diverse Density Concept Demonstrated. Example positive bags (green) and negative bags (red) are embedded into 2D and plotted. A concept capable of differentiating the positive and negative bags is also plotted in the same space. The positive bags have a closer distance to the concept than the negative bags, meaning that there exists a threshold on this distance capable of correctly labeling all bags.

Given that 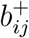 is an instance in bag 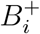, and 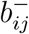 is an instance in bag 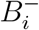, a proposed method to estimate *Pr*(*c*|*B*_*i*_) is the following:

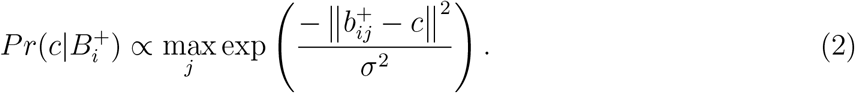

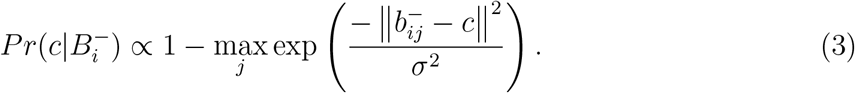

Equation 2 finds the closest distance between any instance in the positive bag 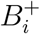 to the concept, maximizing the value towards 1 as the distance decreases. Similarly, Equation 3 finds the same distance, but reaches the maximum value as the distance increases instead. The distance between an instance and the concept is scaled by 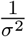. When the scaling factor *σ* is small, an instance needs to be closer to the concept to achieve high similarity.

Multiple-Instance learning via embedded instance selection (MILES) extends the use of a single target concept to the use of many target concepts. With this change, it is now assumed that the ensemble of concepts is capable of representing more complex relationships between the bags than those possible by a single concept that is required to be closer to positive bags. For example, a given concept may instead be closer to the negative bags than to the positive bags. Given a set of concepts *C*, the estimate of *Pr*(*c* |*B*_*i*_) is now defined independently of the bag label:

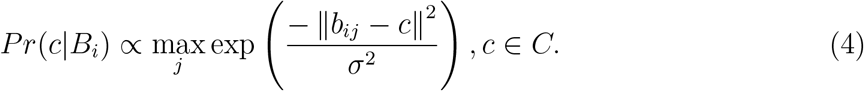

Given *N* = *N* ^+^ + *N* ^−^ where *N* ^+^ is the number of instances of the positive class and *N* ^−^ is the number of instances of the negative class, MILES uses all *N* instances as potential concepts; *C* = {*c*_1_, *c*_2_, …, *c*_*N*}_. If each instance in a bag *B*_*i*_ is represented by a single feature vector *b*_*ij*_, then *B*_*i*_ can be embedded in a *N* -dimension vector *E*_*i*_ through an aggregation of its instances as follows:

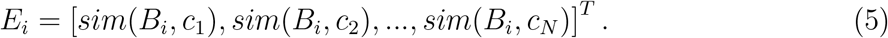

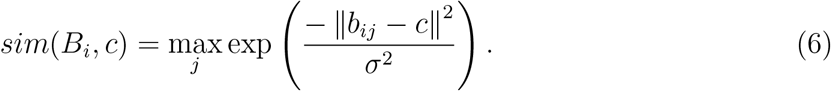

The value of *E*_*ik*_, *k* ∈ {1, 2, …, *N*} is equal to the similarity between the most similar instance in *B*_*i*_ to the concept *c*_*k*_ as defined in Equation 6.

Given all our bags, represented by their embedding vector and a bag label, the algorithm learns to classify the bags through an L1-norm linear support vector machine (SVM) [31]. The SVM learns to fit a hyperplane that best separates the bags based on their label through an optimization process, where the separation is performed in the same space as the bags’ embedding. Since an L1-norm penalty is applied, the optimization process is incentivized to reduce the sum of the absolute value of the hyperplane weights, leading to sparsity. A bag is simply classified on the basis of the side of the SVM hyperplane it resides.

Each dimension in the embedding of a bag is based on the most similar of its instances to a concept, but this concept can be equally similar to multiple instances. For a concept *c*_*k*_, the number of these instances is *m*_*k*_. From the perspective of an instance *b*_*ij*_, there is also a set of concepts to which it was among the most similar in bag *B*_*i*_ to; we denote the set of indices corresponding to these concepts as *I*_*j*_. Given that *w*_*k*_ is the *k*^*th*^ weight in the SVM hyperplane, the classification score of an instance is:

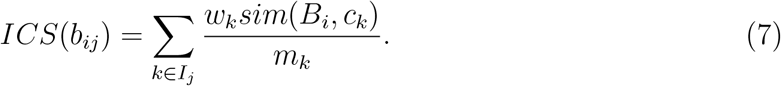

The *ICS* of *b*_*ij*_ is computed by summing all similarity values that *b*_*ij*_ contributed to the bag embedding, with each value scaled by its corresponding SVM weight *w*_*k*_ and normalized by *m*_*k*_, the number of instances equally similar to concept *c*_*k*_. In this step, the required use of a linear SVM is seen, where the weights of the SVM hyperplane can be directly mapped to the features of the bags embeddings. Once the classification score of an instance is calculated, a threshold is used to determine its label. Although the authors of MILES acknowledge that different thresholds may be used, they recommend that anything greater than 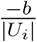 be labeled positive. Where *b* is the SVM hyperplane bias, and *U*_*i*_ is the minimal set of instances in *B*_*i*_ required to create its embedding as defined in Equation 5.

### 2.4 siMILe

Although MILES is powerful, only a single positive instance is theoretically required to correctly classify a bag. This means that a bag containing multiple positive instances can ignore some of them during the creation of the bag embedding or during the SVM learning phase, while still maintaining equivalent performance on bag classification. This susceptibility to using only a fraction of the discriminative features to classify is a problem intrinsic to many such classifiers, hindering MILES’ ability to correctly classify all positive instances. Furthermore, as described, MILES assumes a traditional MIL formulation, making it inefficient when the goal is to compare two classes and extract discriminative instances from each, instead of from only one. For example, as given in Fig. 2B, assume you have classes *A* and *B*, where class *A* contains instances *a* and instances *c*, while class *B* contains instances *b* and instances *c*. When looking to predict the discriminative instances in these classes, which are the instances *a* and *b*, MILES would require two training phases. In the first phase, we would declare class *A* the positive class and label all instances in class *B* negative, since instances *a* are now considered positive instances; MILES would look to label them as positive. During the second run, the classes are swapped and the *B* class is considered positive; the same procedure is used, except with the goal of applying positive labels to instances *b*. This requirement of multiple phases is inefficient and can cause unstable results in the case that the model learns by focusing only on the discriminative instances that exist within the negative class. In this case, positive bags are classified as positive based only on the absence of discriminative instances found in the negative bags, which means that the difference between positive and negative instances in positive bags is not represented in either the bag embedding or the learned SVM weights, to the detriment of instance classification.

To alleviate the problem caused by the use of a subset of discriminative positive instances to classify, we improve MILES by implementing an adversarial erasing scheme (Fig. 4). In adversarial erasing, the goal is to extract all discriminative instances by training the model over multiple iterations and removing the predicted discriminative instances before the next iteration. In the traditional MIL case, this would mean removing positive instances and retraining the classifier until no more positive instances can be found. The iterations will end when the performance of the bag classifier is deemed insufficient. Practically, we set a minimum accuracy in bag classification performance, denoted *minacc*, to determine when to stop further iterations. Adversarial erasing also helps in the situation where the classifier focuses only on the discriminative instances in one class. Should this happen, those instances would be iteratively removed until the classifier is forced to look at the discriminative instances in the other class.

**Figure 4.**
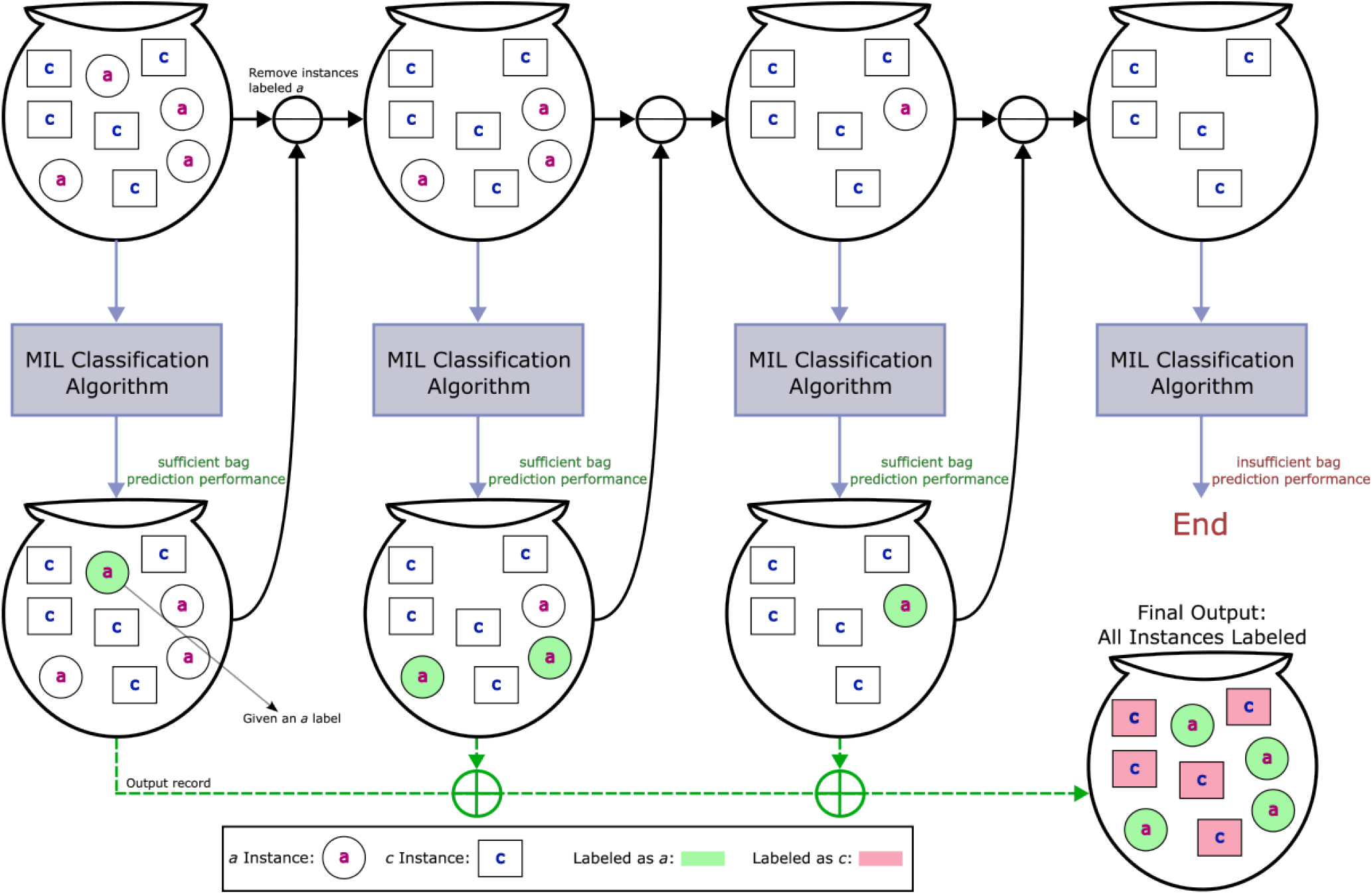
Adversarial erasing applied to multiple instance learning. The visual is given in the context of the formulation where two classes are compared; *A* and *B*, with class *A* having instances *a* and *c*, while class *B* has instances *b* and *c*. The goal in this formulation is to label the instances *a* and instances *b*. The visualization follows the state of a single class *A* bag through adversarial erasing iterations. In each iteration the classifier is trained to separate bags by class, after which it applies labels to the instances in the bag. All instances given a label *a* are removed from the bag before the next iteration. The iterations stop when attempting to train the classifier no longer results in the ability to sufficiently classify bags, at which point all the labels given through the iterations are collected, with the remaining instances labeled as *c*. The instances labeled *a* in early iterations are those that were the focus of the classifier while training at that iteration, by removing them before the next iteration, the classifier is forced to learn based on the remaining.

To optimize the algorithm to classify both classes in a single run, we replace the single threshold classification of the instance classification scores with a k-means scheme using three centers. Given the set of all predicted instances as *X*, a given instance *b*_*ij*_ ∈ *X* is predicted as follows:

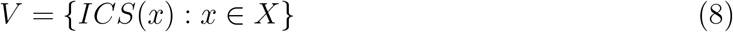

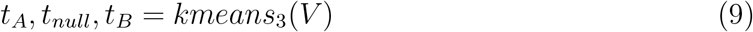

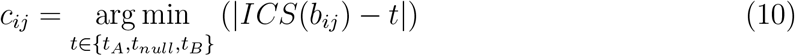

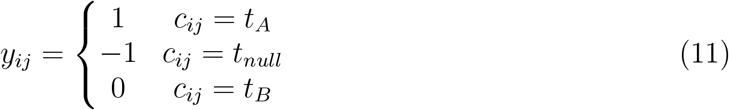

The scores of all predicted instances are aggregated and used with k-means to find three class centers. The label of a given instance is applied based on the center it was closest to. Using this symmetric classifier to classify discriminative instances in both classes, siMILe avoids the computational cost of running the asymmetric base algorithm twice. Furthermore, through the use of clusters, the algorithm is able to be more conservative in its labeling, only labeling instances that cluster around the discriminative centers rather than labeling all instances above a fixed threshold as in traditional MILES. While this could normally be to the detriment of the algorithm’s ability to classify all discriminative instances, the use of adversarial erasing iterations negates this risk and instead enables more precise instance classification (Sec. 2.5).

While it is in principle possible to use deep learning for the classification step, we select an SVM as the basis for training in siMILe to keep training time very short. Although adversarial erasing is powerful in its ability to label more discriminative instances and enable the use of MIL on classes that each contain these discriminative instances, its iterative retraining could become prohibitive when using a deep learning architecture instead. This is particularly seen in cases where these iterations can become numerous. Future work can consider extending this to methods in which retraining can be done using deep learning approaches without retraining from scratch, but this is beyond the scope of this work. By leveraging an SVM, siMILe maintains a relatively low training time, enabling use on less powerful hardware, while also not requiring a graphics processing unit (GPU), and keeping siMILe’s carbon footprint sufficiently small. Our results (Sec 2.5 and onward) show that performance is not meaningfully compromised by these choices.

### 2.5 Evaluation on Simulated Dataset

We first detail the performance of siMILe on simulated data to quantify the effect of our algorithmic improvements between siMILe and MILES. Simulated SMLM data enables us to have full control of the objects and noise model in the data, making it ideal to validate our stated claims that differentiate siMILe from MILES. Using the RSMLM [32] package to simulate dSTORM image acquisition, we generated three types or classes of clustered localizations, or ‘blobs’: A, B, C. We grouped them in two labels or ‘conditions’: Sets with label A containing clusters of type *a* and *c*, while sets with label B contain clusters of type *b* and *c*. (Fig. 5). We compare the ability of siMILe to extract discriminative instances from each class against MILES, MILES with adversarial erasing (MILES + AE), and MILES with the symmetric classifier (MILES + SYM-C). We report the comparison metrics relative to bag size to alleviate the potential issue of specific algorithms having better computational performance based on bag size. The results are also provided based on the average of a nested cross-validation using 5 folds, with the standard deviation reported. The results for F1, precision, recall, and training time are given in Figure 6. Figure 6 shows that siMILe outperforms MILES and the isolated improvements such as SYMC and AE consistently across bag sizes in all metrics. The exception occurs for very small bag sizes, but as the reader can confirm, here all results for all methods are too noisy to extract consistent patterns.

**Figure 5.**
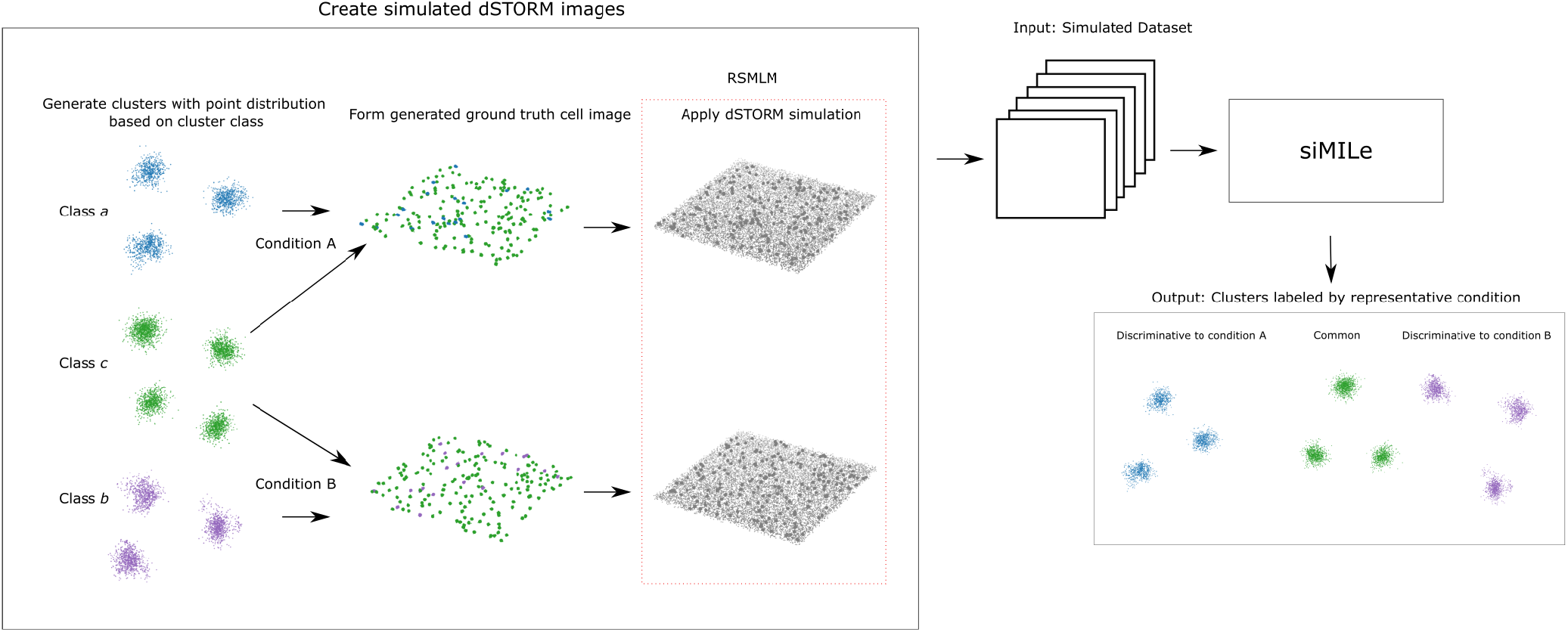
Generating the two conditions of the simulated dSTORM dataset and applying siMILe to identify condition-specific clusters.. The simulated dSTORM dataset consists of 3D point clouds, each labeled as condition *A* or *B*. Condition labels are determined by the cluster classes present within the point clouds, which differ by their generative distributions. Specifically, condition *A* contains clusters from classes *a* and *c*, while condition *B* includes clusters from classes *b* and *c*. These point clouds are processed with the RSMLM [32] package to simulate dSTORM image acquisition. Once multiple datasets for each condition are generated, the siMILe pipeline is applied to identify and label clusters as either condition-specific (discriminative) or common to both conditions..

**Figure 6.**
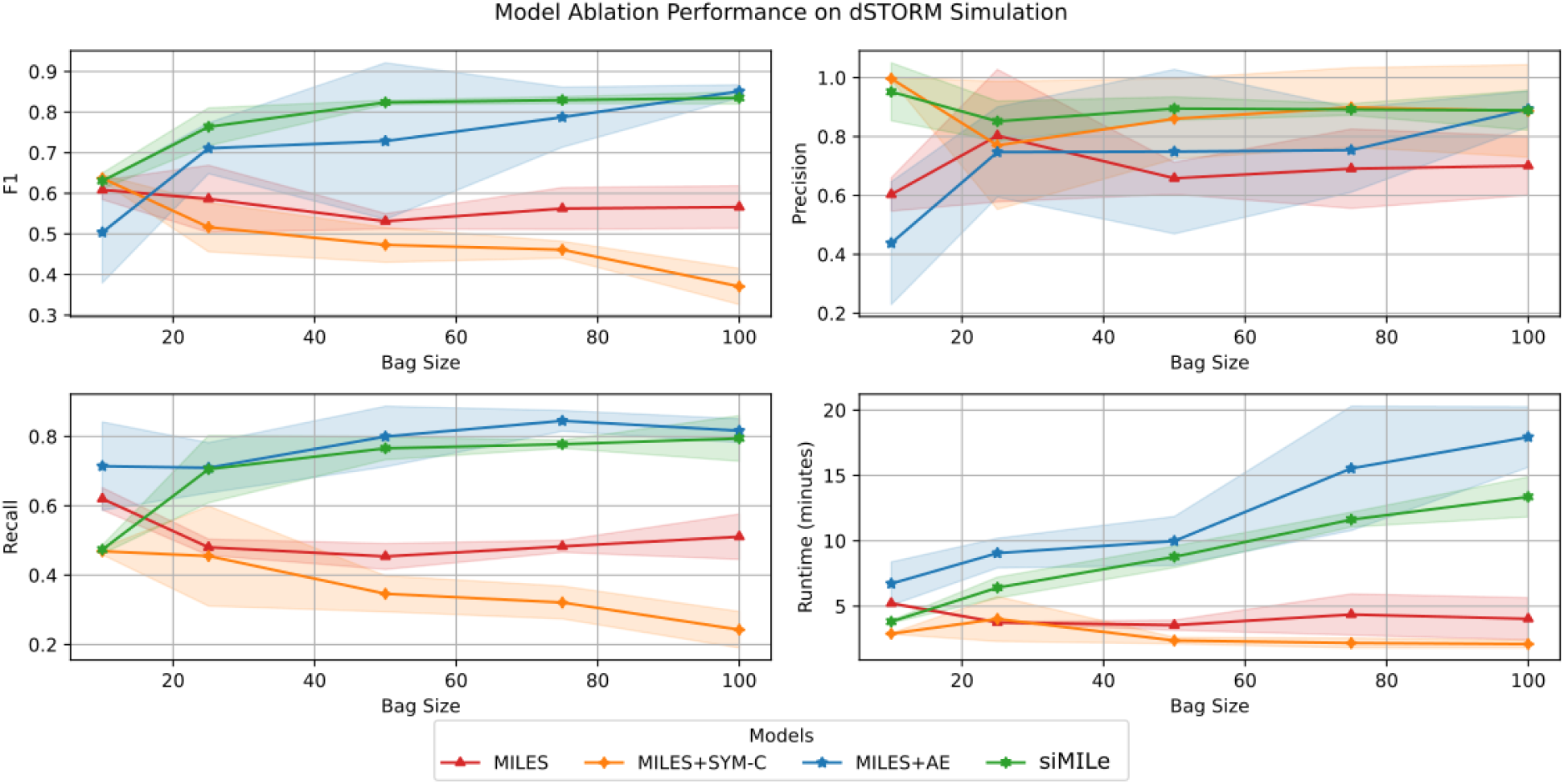
Ablation results on instance classification performance and single core runtime. The highest F1 score is maintained by siMILe across bag sizes, followed closely by MILES + AE, then MILES, with MILES + SYM-C having the worst performance. The recall is similiar to F1 except for MILES + AE performing slightly better than siMILe regardless of bag size. The highest precision scores are siMILe and MILES + SYM-C while MILES has the lowest except as smaller bag sizes. The models using AE have significantly higher training times, increasing with bag size when maximizing classification performance, while MILES and MILES + SYM-C maintain low train times regardless of bag size when achieving their best classification performance. Lines show mean values from nested 5-fold cross-validation with shaded regions indicating ±1 standard deviation.

The training time of a model is recorded using an Intel Core i7-7740X with 32GB RAM available on a machine running Ubuntu 20.04; the time is reported based on use of a single core. Models using AE include all iterations in their recorded runtime. Runtime results show that siMILe has quite short training times (*<* 20 min), enabling users to retrain it quickly if needed. siMILe has an increased training time compared to MILES, trading training time for improved performance. Given the low overall training time, this is a worthwhile compromise.

### 2.6 siMILe identifies Caveolae as Discriminative Structures between PC3 and PC3-CAVIN1 cells

Next, we evaluate siMILe on its capability to detect complex discriminative protein structures in real dSTORM data using the PC3/PC3-CAVIN1 dataset [9]. Considering that Cav1 does not form caveolae in the absence of cavin-1, we investigate the ability of siMILe to detect caveolae in cavin-1-expressing PC3 cells (referred to as PC3-CAVIN1) in comparison to PC3 cells lacking cavin-1 and not presenting caveolae [33].

Using the features produced by SuperResNET, we apply siMILe to classify the clusters in PC3-CAVIN1 as discriminative to PC3-CAVIN1 or common to PC3 (not discriminative) (Fig. 7). Our results labeled 8% of blobs within PC3-CAVIN1 as discriminative (Fig. 8). Blobs labeled discriminative to PC3-CAVIN1 cells are more spherical and much larger than the common blobs. SuperResNET classification (which categorizes Cav1 structures as S1A scaffolds/8S complexes, S1B and S2 scaffolds as intermediate oligomers, or caveolae based on size and morphology) of the discriminative blobs shows that they consist predominantly of caveolae, with a smaller proportion of S2 scaffolds, 1 S1B scaffold and no S1A scaffolds (Fig. 8B). This is consistent with the 200-250 nm size and 90 nm distance to centroid of the point clouds, as well as localization counts approaching that of the estimated 144±39 in caveolae [34]. It is clear that the model is identifying caveolae-like structures as discriminative to PC3-CAVIN1.

**Figure 7.**
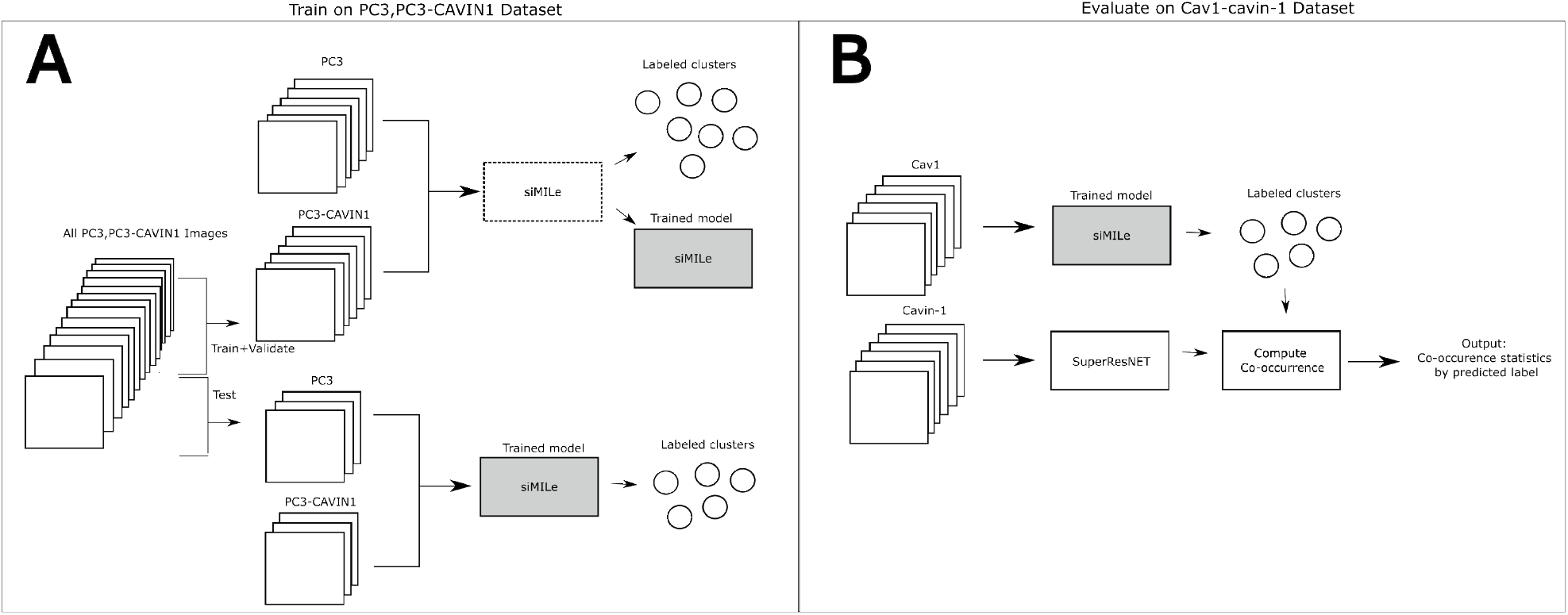
Experiment setup for the application of siMILe to PC3,PC3-CAVIN1 and two-channel Cav1-cavin-1 datasets. (A) The PC3,PC3-CAVIN1 3D point cloud dataset is split into training, validation, and testing subsets. The training and validation sets are used to train siMILe and fine-tune hyperparameters for labeling clusters. The final trained model is applied to the test set to generate the reported results. (B) The pre-trained model from A is applied to the Cav1 channel of the Cav1-cavin-1 dataset. Clusters identified by the model are labeled and compared against the SuperResNET-processed cavin-1 channel to assess co-occurrence by label.

**Figure 8.**
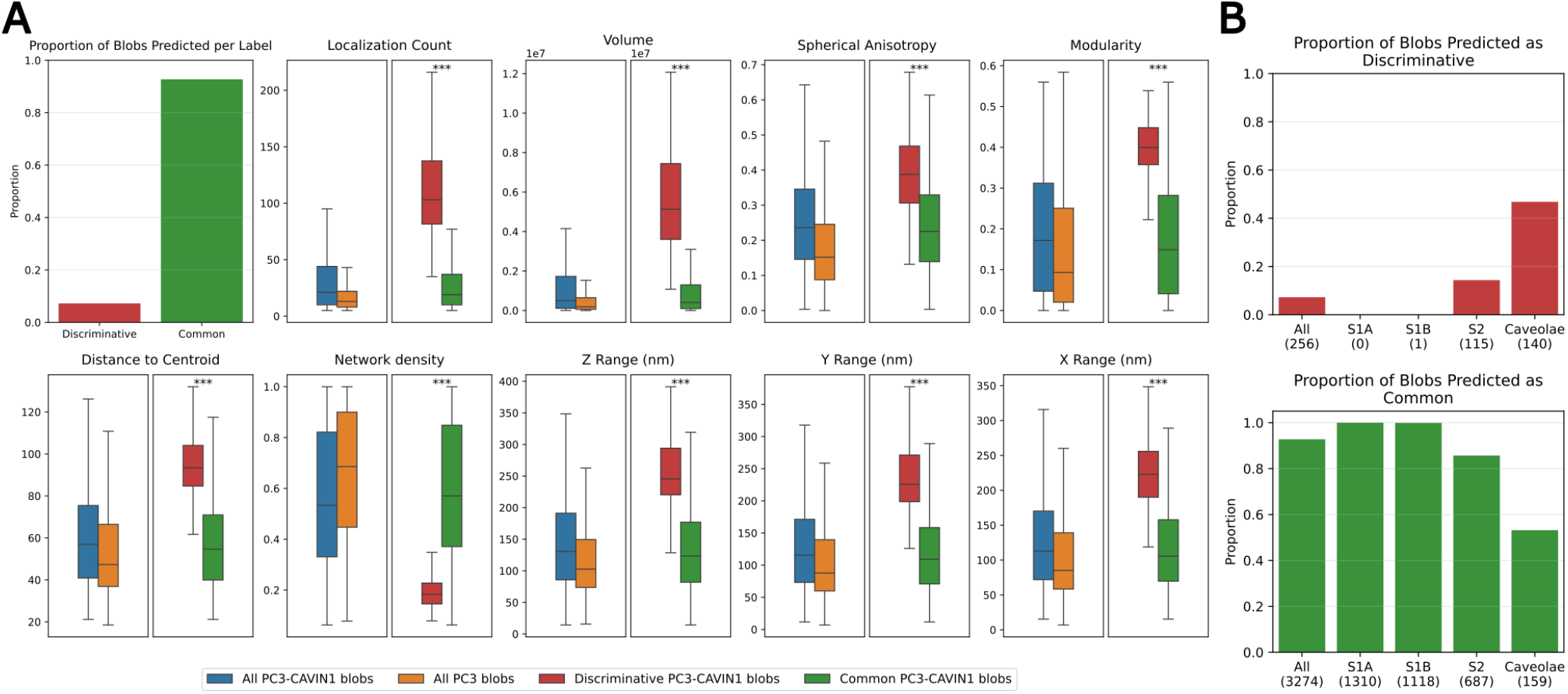
Feature comparison of predicted blob labels in the PC3,PC3-CAVIN1 dataset. After applying siMILe to compare the PC3 and PC3-CAVIN1 conditions, we show the proportion of labels predicted in the first plot out of the 7060 blobs, with ∼93% of the blobs labeled as common. The feature distribution of these blobs, based on the predicted labels (discriminative in red and common in green), is compared against the feature distribution of all blobs in the PC3 (orange) and PC3-CAVIN1 (blue) datasets. The discriminative blobs are consistently larger in size with many more localizations. They also appear to be more spherical, contain less dense networks, and contain more modules. These features are signature for caveolae, one of the known unique differences between PC3 and PC3-CAVIN1. This shows that siMILe is able to isolate caveolae as unique to PC3-CAVIN1, given that they require cavin-1 to form. Objects identified as caveolae in panel B are classified using SuperResNET. It is possible that SuperResNET mislabels some S2 as caveolae and vice versa, which can explain the partial agreement of siMILe and SuperResNET. Box plots show median (center line), interquartile range (box), and 1.5× IQR (whiskers). Statistical comparisons were performed using the two-sided Mann-Whitney U test. Significance levels: ^∗^p *<* 0.05, ^∗∗^p *<* 0.01, ^∗∗∗^p *<* 0.001. Non-significant comparisons (p ≥ 0.05) are not marked.

### 2.7 Transfer of Trained Model to Dual-Channel Cav1-cavin-1 Dataset

Having identified discriminative structures in PC3-CAVIN1 cells, we next sought to validate these findings using an independent dataset with biological ground truth. We applied the trained model to PC3-CAVIN1 cells dually labeled for both Cav1 and cavin-1, where cavin-1 co-localization provides independent validation of caveolar structures. The Cav1 channel was processed by SuperResNET to identify S1A, S1B and S2 scaffolds as well as caveolae [9] and then overlaid with the cavin-1 labeled channel. Representative two-channel dSTORM images show the spatial distribution of Cav1 and cavin-1 in PC3CAVIN1 cells, with SuperResNET classification revealing the organization of different Cav1 structural classes relative to cavin-1 (Fig. 9).

**Figure 9.**
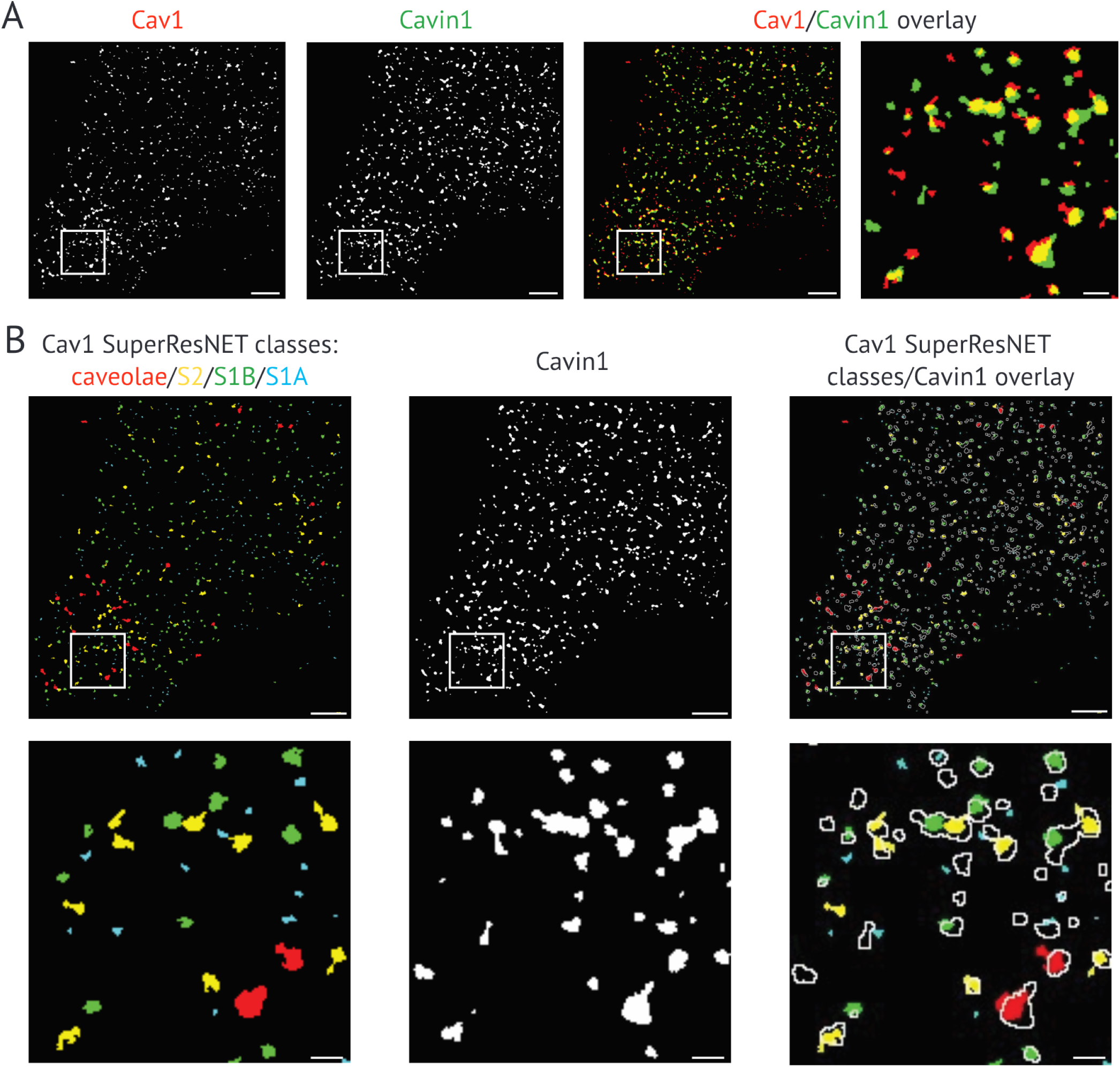
SuperResNET classification of two-channel Cav1-cavin-1 dSTORM images (A) 2D mask overlay of 3D Cav1-cavin-1 two-channel dSTORM images. Scale bars: 2 µm (whole image) and 150 nm (inset). (B) Overlay of Cav1 SuperResNET classes (red: caveolae, yellow: S2, green: S1B, cyan: S1A) and cavin-1. Scale bars: 3 µm (whole image) and 250 nm (inset).

Application of the PC3/PC3-CAVIN1 trained model to the Cav1 channel of this new dataset identified 22% of blobs as discriminative (Fig. 10A). Feature distributions of discriminative versus common blobs showed similar patterns to those observed in the original dataset, with discriminative blobs exhibiting larger size, higher localization counts, increased sphericity, less dense networks, and more modules. The normalized Euclidean distance between blob classes in both datasets confirmed high feature similarity (Fig. 10B), demonstrating successful model transfer.

**Figure 10.**
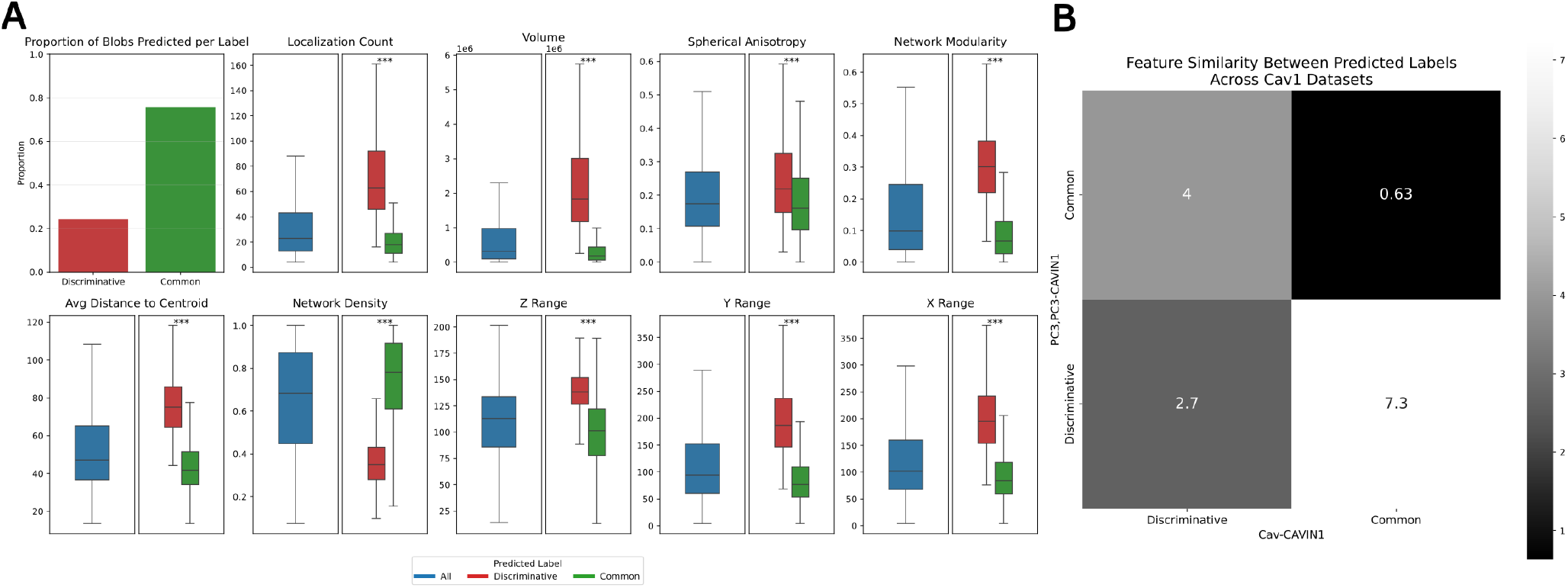
Feature comparison of predicted blob labels in the Cav1-cavin-1 dataset from trained model. (A) After applying siMILe trained on PC3,PC3-CAVIN1 to the Cav1 channel of the Cav1-cavin-1 dataset, the proportion of labels is shown in the first plot, with the majority of the 30,412 blobs labeled as common. The feature distribution of these blobs, based on the predicted labels (discriminative in red and common in green), is compared against the feature distribution of all blobs in the Cav1 (blue) images. The discriminative blobs consistently follow the trend seen in the PC3-CAVIN1 predictions. These blobs are of a larger size, contain a higher count of localizations, are more spherical, have less dense networks, and have more modules. (B) The predicted labels are compared between both datasets through a normalized euclidean distance, supporting the similarity of these labels across both datasets. Box plots show median (center line), interquartile range (box), and 1.5× IQR (whiskers). Statistical comparisons were performed using the two-sided Mann-Whitney U test. Significance levels: ^∗^p *<* 0.05, ^∗∗^p *<* 0.01, ^∗∗∗^p *<* 0.001. Non-significant comparisons (p ≥ 0.05) are not marked.

### 2.8 Cavin-1 Interaction Validates Discriminative Structure Identification

To determine whether discriminative structures identified by siMILe represent biologically relevant populations, we assessed their association with cavin-1. To quantify the spatial proximity between Cav1 and cavin-1 structures, we calculated a Blob Overlap Parameter (BOP) for each Cav1 blob as − log(*d/*(*r*_cav1_ + *r*_cavin-1_)), where *d* is the distance between the Cav1 blob centroid and its nearest cavin-1 blob centroid, and *r* represents the respective radii. Thus, a BOP value of 0 indicates structures with touching borders, positive values indicate overlapping structures with higher values representing greater overlap, and negative values indicate separated structures with more negative values representing greater separation.

Correlation analysis revealed that features associated with discriminative labeling also correlated with cavin-1 interaction (Fig. 11A, all correlations p*<*0.001). Network density showed the strongest correlation product (0.32), with both discriminative and cavin-1-interacting structures exhibiting lower density values. Distance to centroid metrics all correlated positively with both parameters, yielding correlation products of 0.30 (average), 0.29 (median), and 0.28 (maximum). Spatial parameters (Y range: 0.28, X range: 0.27), structural features (max degree: 0.28, area: 0.27), and network modularity (0.23) also showed positive correlations with both discriminative labeling and cavin-1 interaction. Characteristic path length showed negative correlations with both parameters (correlation product: 0.27).

**Figure 11.**
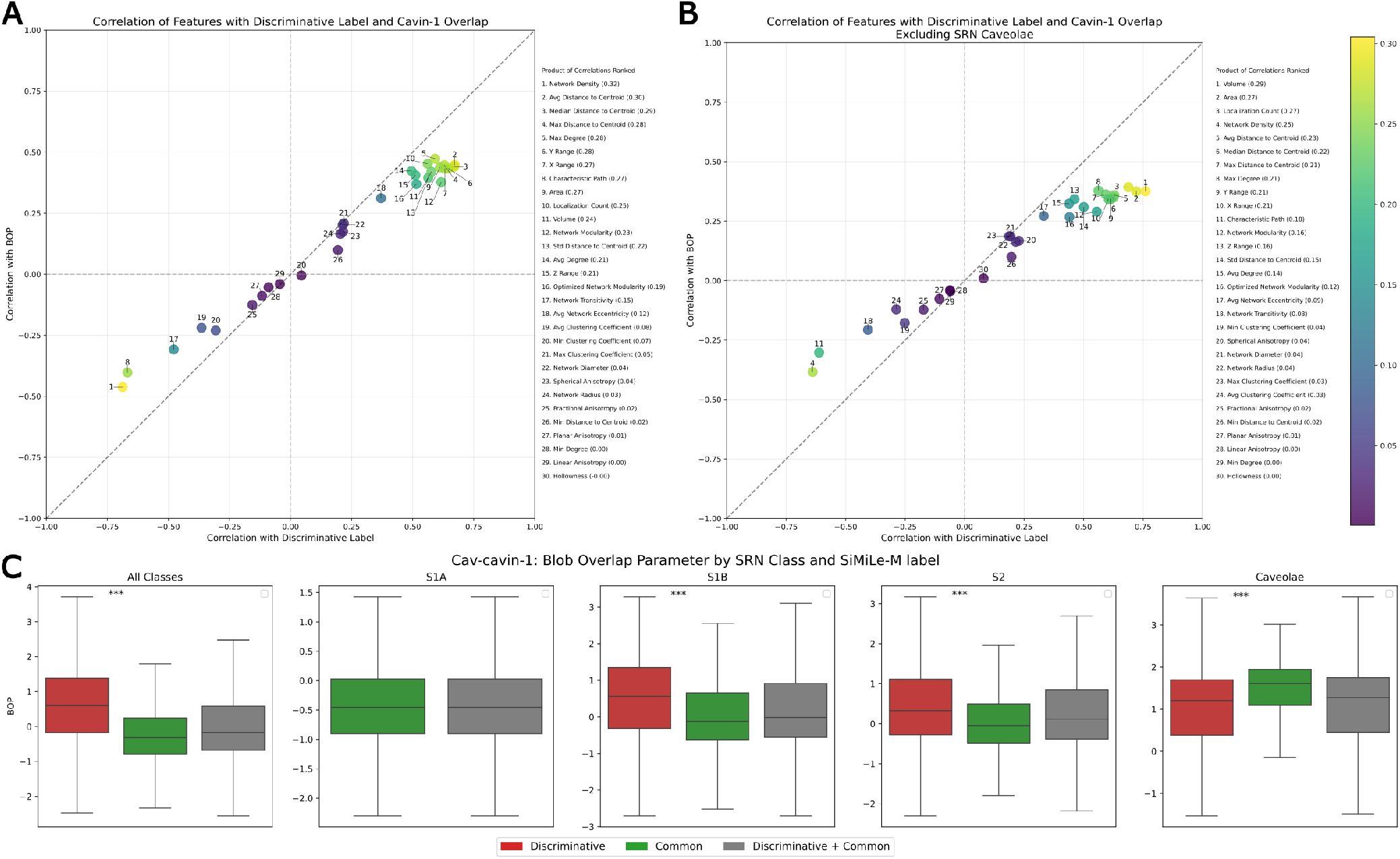
Correlation analysis of network features with discriminative labels and cavin-1 interaction. (A) Scatter plot showing the correlation of blob features with the discriminative label (x-axis) and cavin-1 interaction (y-axis). Features follow a positive correlation trend, indicating a consistent relationship between discriminative properties and increased cavin-1 interaction. The product of correlations ranked (right) highlights network density, distances to centroid, and spatial range parameters as top contributors. (B) Similar correlation analysis excluding SRN caveolae samples. The exclusion slightly alters correlation rankings, with volume, area, localization count, and network density being the strongest contributors. (C) Box plots displaying cavin-1 Blob Overlap Parameter (BOP) distributions across different SRN classes (All Classes, S1A, S1B, S2, and caveolae), separated by siMILe predicted labels (discriminative in red, common in green, combined in gray). Across SRN S2 and S1B, discriminative blobs exhibit larger overlap compared to common blobs based on the larger BOP values, indicating that siMILe identifies structures with cavin-1 association patterns despite not being SRN caveolae. Box plots show median (center line), interquartile range (box), and 1.5× IQR (whiskers). Statistical comparisons were performed using the two-sided Mann-Whitney U test. Significance levels: ^∗^p *<* 0.05, ^∗∗^p *<* 0.01, ^∗∗∗^p *<* 0.001. Non-significant comparisons (p ≥ 0.05) are not marked.

Analysis excluding SuperResNET-classified caveolae showed similar correlation patterns for scaffold structures (Fig. 11B, all correlations p*<*0.001). Size-related features showed the highest correlation products, with volume (0.29), area (0.27), and localization count (0.27) all positively correlating with both discriminative labeling and cavin-1 interaction. Network density maintained negative correlations with both parameters (0.25), while distance to centroid metrics showed positive correlations (0.21-0.23). Network modularity showed positive correlations (0.16), though reduced compared to the full dataset.

Examination of BOP across SuperResNET classes showed discriminative blobs had higher BOP values than common blobs within each SRN class (Fig. 11C). This difference was most pronounced for S1B and S2 scaffolds, while caveolae showed similar BOP for both populations. SuperResNET classification analysis showed 86% of caveolae were identified as discriminative, compared to 55% of S2 scaffolds, 26% of S1B scaffolds, and 0% of S1A scaffolds (Fig. 12). Discriminative caveolae were smaller with fewer localizations than common caveolae. Discriminative S1B and S2 scaffolds showed lower network density, higher distance to centroid values, larger spatial ranges, and greater size metrics compared to common structures, with network modularity differences most pronounced in S1B.

**Figure 12.**
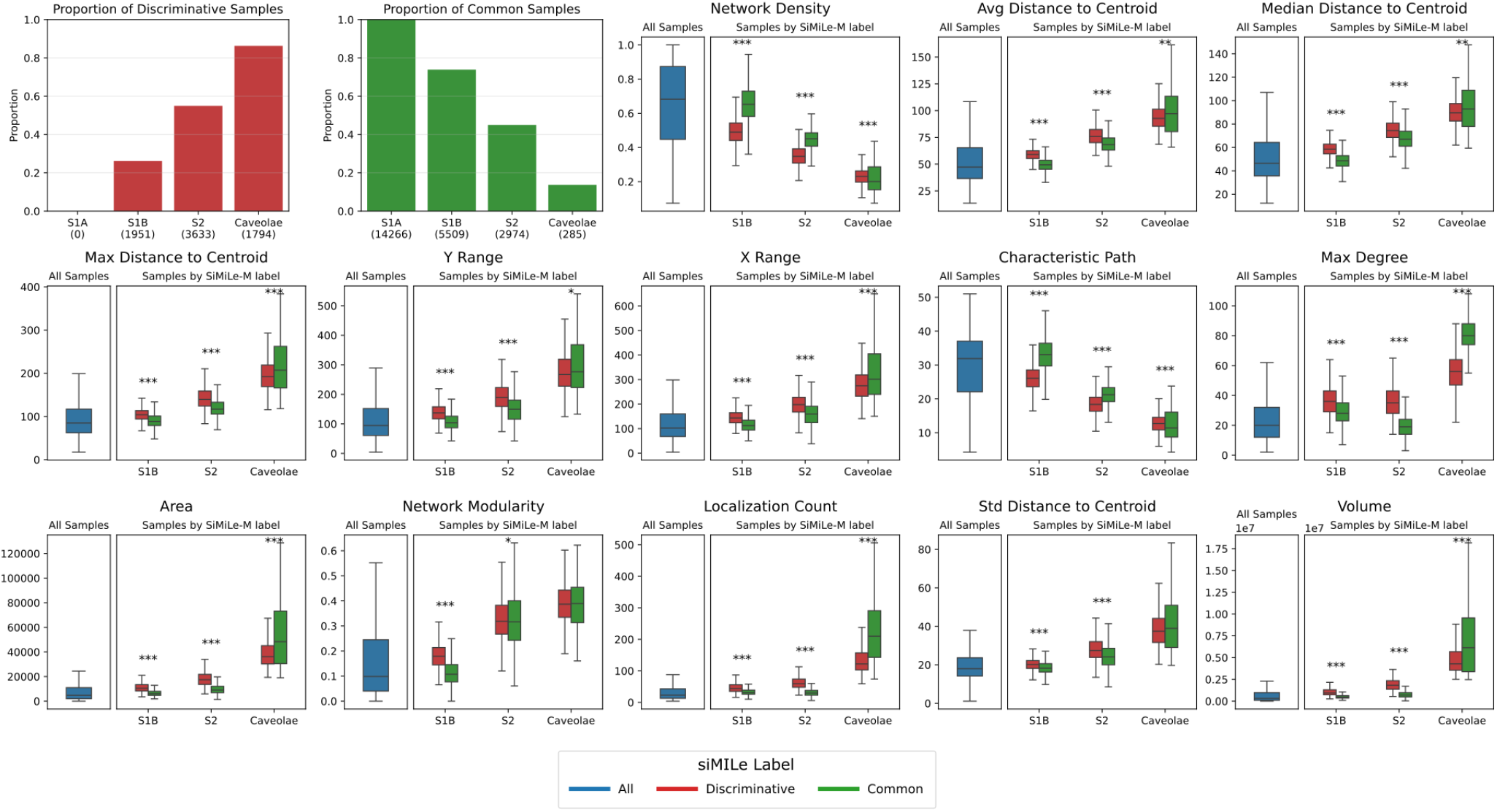
Feature comparison of discriminative and common blobs within SuperResNET classes. Results from applying the PC3/PC3-CAVIN1 trained model to the Cav1-cavin-1 dataset (30,412 blobs), displayed as a 3×5 grid. First two panels show the proportion of discriminative and common labels within each SRN class (S1A: 0% discriminative; S1B: 26%; S2: 55%; caveolae: 86%). Remaining panels show feature distributions for all samples (blue), as well as discriminative (red) versus common (green) blobs within each class. For S1B and S2 scaffolds, discriminative blobs exhibit lower network density, higher distance to centroid statistics, larger spatial ranges, and greater size metrics, with network modularity differences very pronounced in S1B. Common caveolae show larger values across size-related features than discriminative caveolae. S1A scaffolds excluded due to absence of discriminative blobs. Box plots show median (center line), interquartile range (box), and 1.5× IQR (whiskers). Statistical comparisons were performed using the two-sided Mann-Whitney U test. Significance levels: ^∗^p *<* 0.05, ^∗∗^p *<* 0.01, ^∗∗∗^p *<* 0.001. Non-significant comparisons (p ≥ 0.05) are not marked.

These results demonstrate that siMILe, using only weak supervision, identifies structures with enhanced cavin-1 interaction that include not only the expected caveolae but also S1B and S2 scaffold populations.

### 2.9 Identification of Discriminative Clathrin-Coated Pits Upon Small Molecule Inhibition

To demonstrate the flexibility and generality of siMILe, we applied siMILe to the previously published SMLM dataset clathrin-coated pits imaged by labeling the clathrin heavy chain in HeLa cells treated with the clathrin endocytosis inhibitors pitstop 2, dynasore or latrunculin A (LatA) [35]. See summary of data analyzed in Table 1.

**Table 1.** Summary of the clathrin endocytosis dataset from Wong et al. [35].

| Compared Conditions |  | Number of cells<br>in each replicates<br>(=Total) | Number<br>of training<br>samples | Number of<br>Validation<br>samples | Number<br>of Testing<br>samples | Total<br>number |
| --- | --- | --- | --- | --- | --- | --- |
| DMSO vs Pitstop 2 | DMSO | 6 + 7 + 8 + 6 +<br>7 + 6 (=40) | 11363 | 2418 | 2176 | 15957 |
|  | Pitstop 2 | 7 + 7 + 6 + 6 +<br>6 (=32) | 6529 | 1526 | 2647 | 10702 |
| DMSO vs LatA | DMSO | 6 + 7 + 8 + 6 +<br>7 + 6 (=40) | 11363 | 2418 | 2176 | 15957 |
|  | LatA | 7 + 7 + 7 (=21) | 1250 | 850 | 1280 | 3380 |
| DMSO vs Dynasore | DMSO | 6 + 7 + 8 + 6 +<br>7 + 6 (=40) | 11363 | 2418 | 2176 | 15957 |
|  | Dynasore | 7 + 7 + 7 (=21) | 3529 | 2474 | 2945 | 8948 |

Clathrin-mediated endocytosis proceeds through various stages, in which cargo binding initiates the formation of a clathrin-coated pit that matures, resulting in membrane invagination and dynamin-dependent scission of the vesicle from the plasma membrane [36]. Small molecule inhibitors of clathrin-mediated endocytosis include pitstop 2, dynasore and the actin depolymerizing agent LatA [37, 38]. Pitstop 2 binds directly to the N-terminal domain of the clathrin heavy chain [39], dynasore is a non-competitive inhibitor of dynamin, required for clathrin vesicle scission [40, 41], while latrunculin A (LatA) directly binds to actin monomers to inhibit F-actin polymerization inhibiting clathrin endocytosis [38]. Prior application of SuperResNET analysis to clathrin-coated pits imaged by labeling the clathrin heavy chain in HeLa cells identified clathrin structures corresponding to clathrin pits and vesicles (class II) [35]. SuperResNET classification feature analysis showed that dynasore and pitstop 2 reduced the size of clathrin pits and vesicles compared to control cells with pitstop 2 more significantly reducing the size of clathrin pits and vesicles compared to dynasore-treated pits.

Here, we applied siMILe to extract discriminative blobs in pitstop 2, dynasore and LatA treated cells compared to untreated HeLa cells. siMILe detects a higher percentage of discriminative blobs for pitstop 2 compared to either dynasore or LatA (Fig. 13A). This suggests that pitstop 2 causes a more distinct structural shift in clathrin pits, perhaps related to the fact that pitstop 2 binds directly to the clathrin heavy chain [39]. Consistent with SuperResNET feature analysis [35], discriminative blobs identified by siMILe upon pit-stop 2 and dynasore inhibition result in blobs with smaller size features compared to control, while LatA blobs are larger with higher spherical anisotropy (Fig. 13B). Comparison of discriminative pitstop 2 and dynasore class II blobs shows that dynasore blobs are distinctly larger in their x, y and z ranges, have a higher localization count, and are less linear and more spherical (Fig. 13C). These features suggest that discriminative dynasore blobs are further along the process of clathrin pit maturation than pitstop 2 blobs, as suggested previously [35]. siMILe is therefore able to detect structural differences to clathrin-coated pits induced by small molecule inhibitors.

**Figure 13.**
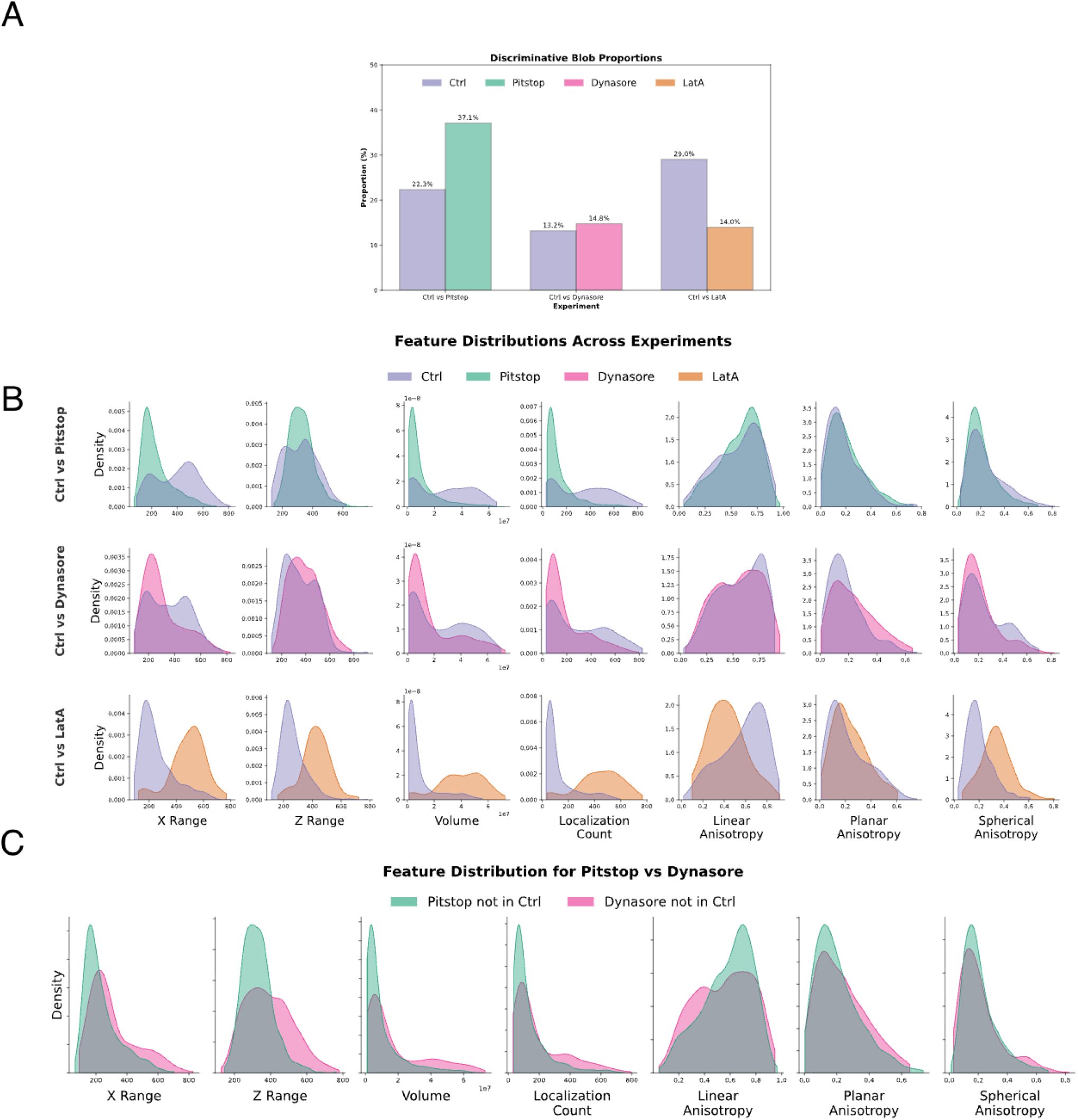
Applying siMILe to clathrin endocytosis datasets to contrast different treatments: Pitstop, Dynasore, and LatA to the control (Ctrl) condition. The SMLM datasets are analyzed with SuperResNET to extract features of clathrin pits (blobs or clusters). (A) The proportions of discriminative blobs when siMILe was used to compare Ctrl to Pitstop; Ctrl to Dynasore, and Ctrl to LatA. (B) The density distribution of the blob features for the same 3 siMILe experiments. (C) Comparing the feature distribution between blobs that are discriminative to Pitstop (vs Ctrl) with the ones that are discriminative to Dynasore (vs Ctrl).

## 3 Discussion

### 3.1 siMILe: an improved MIL approach for SMLM

We introduce siMILe, an enhanced MIL approach incorporating MILES with adversarial erasing, to identify, using weak-supervised learning only, differences in protein oligomer structures in cells in SMLM. Application of siMILe to SuperResNET-processed 3D SMLM point cloud data effectively identifies caveolae that are selectively found in PC3 cells expressing cavin-1 [9, 33]. We validate siMILe using simulated and real data, and confirm our findings using known interaction of a second protein (cavin-1) to illustrate that siMILe detects differential structures with biological basis. The use of MIL to label individual structures in 3D point cloud SMLM data has yet to be explored. Our simulation results support the ability of siMILe to improve the classification of more discriminative instances through its very high recall and the significant increase in MILES recall with the addition of adversarial erasing (Fig. 5). The combination of the symmetric classifier with adversarial erasing allows siMILe to maintain very high precision through many iterations that can conservatively label. While MILES can accurately identify discriminative structures when analyzing larger datasets (high precision), it tends to miss many actual discriminative structures (low recall) - a limitation that siMILe overcomes through its combined approach.

Overall, the consistently high F1 score regardless of bag size supports siMILe improvements over MILES. The computational load of siMILe increased significantly with adversarial erasing when maximizing classification performance. We do however, observe, a reduction in this time based on the use of the symmetric classifier. Finally, across all metrics, the standard deviations of siMILe results are very low, almost always the lowest. This indicates that the contributions cause improved stability in siMILe, alleviating sensitivity to parameters. Computation time is longer as the bag size increases when using adversarial erasing. This shows that only a few discriminative instances are required to perform bag classification. As the bag size gets larger, it takes more iterations to remove all discriminative instances, whereas smaller bag sizes allow more of these instances to be reflected in their bag embedding and during training, leading to correct labeling of them during the instance classification step. It could be claimed that a different solution to this problem would be to use only small bag sizes instead of adversarial erasing. The issue with this approach is that it is only possible when the data is well understood and the witness rate known; otherwise, it cannot be determined which bag size contains few enough discriminative instances. This requirement is reasonable in some applications, but not always possible in discovery.

### 3.2 Application of siMILe to SMLM Datasets

SuperResNET previously selectively identified caveolae in PC3 prostate cancer cells transfected with cavin-1 [9]. Using the same dataset we now show that siMILe identifies caveolae as discriminative structures in PC3-CAVIN1 cells demonstrating its ability to detect known structures unique to cavin-1 transfected PC3 cells. siMILe also detected higher order S1B and S2 scaffolds as distinct but not isolated 8S complexes. The 8S complex is a highly stable, SDS-resistant grouping of 11 Cav1’s that exhibits a barrel-like shape by cryoEM [42, 43]. The 8S complex was identified as S1A scaffolds by SuperResNET and shown to be modules that combine to form higher order scaffolds and caveolae [11]. That no S1A complexes were found to be unique to PC3-CAVIN1 cells highlights the stable structure of the 8S complex and the robust ability of siMILe to selectively detect discriminatory structures between datasets.

Extension of the approach to a new dataset in which PC3-CAVIN1 cells are labeled for both Cav1 and cavin-1 shows that siMILe effectively selects cavin-1 labeled structures. These include caveolae but also higher order S1B and S2 scaffolds but not 8S complexes (S1A scaffolds). Caveolae show a close association with cavin-1 with siMILe identifying 80% of SuperResNET defined caveolae as discriminatory. Feature analysis highlights the larger size of common caveolae structures, suggesting that these might be closely associated, overlapping and unsegmented caveolae that are classed by SuperResNET as caveolae based on size features. Discriminatory S1B and S2 scaffolds were found to more closely associate with cavin-1 than common S1B and S2 and present larger and more spherical shape features (Fig. 12). This suggests that cavin-1 association with these oligomeric 8S complexes impacts their structure and their organization.

Cavin-1 is thought to selectively associated with large 70S Cav1 oligomers at the plasma membrane [33, 44]. Our data show that cavin-1 can associate with higher order 8S oligomers, localized to the plasma membrane by TIRF microscopy, and suggests that progressive association of cavin-1 with 8S complex oligomers contributes to caveolae formation. siMILe therefore identifies caveolae from cavin-1 expressing PC3 cells more effectively than SuperResNET and identifies distinct conformations of 8S complex oligomers that are associated with cavin-1.

In addition, we applied siMILe to a clathrin heavy chain SMLM data set treated with clathrin endocytosis inhibitors [35]. siMILe effectively identified smaller clathrin pits and vesicles induced by treatment with pitstop 2 and dynasore, that prevent clathrin pit maturation and vesicle scission, respectively, and larger ones induced by treatment with the actin depolymerization agent LatA, as previously reported [35, 38, 40].

siMILe is a novel MIL-based algorithm that incorporates MILES and adversarial erasing to identify discriminatory structures based on SuperResNET SMLM data analysis. siMILe is therefore able to discover novel changes in protein oligomer structure conditional on cell type, genomic, or environmental labels. Uniquely, siMILe is designed to tackle multiple labels without compromising scalability, and remains interpretable.

## 4 Conclusion

We introduced siMILe, a novel enhancement on the MIL paradigm, to identify, using weak-supervised learning only, differences in protein oligomer structures in cells in SMLM. We validate siMILe using simulated and real data, and confirm our findings using known interaction of a second protein (cavin-1) to illustrate the siMILe detects differential structures with biological basis. siMILe will open the door to novel discovery of changes in protein oligomer structure conditional on cell type, genomic, or environmental labels. Uniquely, siMILe is through its symmetric classifier better positioned to tackle multiple labels without compromising scalability, and remains interpretable when applied to interpretable features.

## 5 Materials and methods

### 5.1 PC3, PC3-CAVIN1 Dataset

A previously published SMLM image dataset of human PC3 prostate cancer cells is used [9]. Although PC3 prostate cancer cells express Cav1 (CAV1; UniProtID: Q03135), they do not express cavin-1 (CAVIN1; UniProtID: Q6NZI2), producing no caveolae. Through stable transfection of cavin-1 in PC3 cells, caveolae can be induced [33]. The dataset contains the PC3 cells absent of cavin-1/PTRF (referred to as PC3) and PC3 cells transfected with cavin-1/PTRF (referred to as PC3-CAVIN1 cells). The images are acquired over three replicates, each replicate contains 10 to 11 images; for both PC3 and PC3-CAVIN1 cells.

### 5.2 Simulated dSTORM dataset

We generate a simulated dSTORM point cloud dataset for testing siMILe on data with ground truth. The point clouds were generated with the RSMLM software package, used in previous publications to test clustering methods in simulations [45, 46], and the dSTORM simulation parameters were adopted from [46]. We generate the dataset with two classes, *A* and *B*, using the setup visualized in Figure 2B. Each class contains clusters representing the structure instances that belong to one of the two classes. The class *A* contains cluster labels *a* and *c*, while the class *B* contains clusters labeled *b* and *c*. The centroid of each cluster is uniformly distributed. The number of localizations is generated using a normal distribution with a mean of 50 and standard deviation of 10. The position of the cluster localizations is also generated with a normal distribution around the centroid, with the *c* clusters generated with equal standard deviation in *x, y, z* of 20, whereas the *a* and *b* instances differ in their *x* and *z* standard deviation, respectively, set at 40. For each class there were 50 cells generated with ∼200 blobs per cell and a witness rate of 10%.

### 5.3 Dual-Channel Cav1-cavin-1 Dataset

#### 5.3.1 Antibodies and Plasmid

Rabbit anti-Cav1 (#3267) was purchased from Cell Signaling and mouse anti-GFP (A11120) was purchased from Invitrogen. Secondary goat anti-rabbit Alexa Fluor 647 F(ab)’2 (A-21246) was purchased from Invitrogen, and goat anti-mouse CF 568 F(ab)’2 (20109) was purchased from Biotium. PTRF/cavin-1-EGFP plasmid was a generous gift from Dr. Michelle Hill (The University of Queensland Diamantina Institute, Brisbane, Australia) [33].

#### 5.3.2 Cell Culture and Transfection

The PC3 cell (RRID: CVCL 0035) clonal line described in previous study [47] was maintained at 37 °C, 5% CO2 in RPMI-1640 medium (Thermo-Fisher Scientific Inc.) supplemented with 10% fetal bovine serum (FBS, Thermo-Fisher Scientific Inc.) and 2mM L-glutamine (Thermo-Fisher Scientific Inc.). The cells were passaged using 0.25% Trypsin-EDTA (Thermo-Fisher Scientific Inc.) at approximately 70% confluency and were discarded at the 10th passage. The cells were tested regularly for mycoplasma using a PCR kit (Catalogue# G238; Applied Biomaterial, Vancouver, BC, Canada)Joshi, 2008. Plasmid transfection was done 24 hours after seeding the cells using Lipofectamine 2000 (Life Technologies, Thermo Fisher Scientific) following manufacturer’s protocol.

#### 5.3.3 SMLM Preparation and Imaging

Coverslips (No. 1.5 H) were sonicated for 30 minutes in 1M potassium hydroxide followed by 30-minute sonication in 100% ethanol and then rinsed with Milli-Q water [12]. The cells were fixed 18 hours after transfection using 4% paraformaldehyde (PFA) in phosphate-buffered saline containing 1 mM MgCl2 and 0.1 mM CaCl2 (PBS-CM) for 15 minutes, rinsed thrice with PBS-CM, permeabilized using 0.2% Triton X-100 diluted in PBS-CM, incubated with Image-iT FX Signal Enhancer (Thermo Fisher Scientific) and blocked using BlockAid Blocking Solution (Thermo Fisher Scientific) [12]. The cells were incubated with primary rabbit anti-Cav1 and mouse anti-GFP diluted in saline sodium citrate (SSC) buffer containing 1% BSA, 2% goat serum and 0.05% Triton X-100 overnight at 4 °C and then with secondary F(ab’)2-goat anti-rabbit Alexa Fluor 647 and F(ab’)2-goat anti-mouse CF568. Cells were washed with SSC buffer containing 0.05% Triton X-100 and post-fixed with 4% PFA for 15 minutes. The cells were then incubated with 0.1 *µ*m TetraSpeck Fluorescent Microspheres (Thermo Fisher Scientific) overnight. Before imaging, samples were mounted in freshly prepared blinking buffer containing 10% glucose (Sigma-Aldrich Inc.), 0.5 mg/ml glucose oxidase (Sigma-Aldrich Inc.), 40 *µ*g/mL catalase (Sigma-Aldrich Inc.), 50 mM Tris, 10 mM NaCl and 50 mM *β*-mercaptoethanol (*β*ME; Sigma-Aldrich Inc.) in Milli-Q water and sealed on glass depression slide. Three replicates of dSTORM images (at least 10 randomly selected images per replicate) were acquired using a Leica SR GSD 3D system with a 160 × objective lens (HC PL APO 160 × /1.43, oil immersion), a 642 nm laser line, a 542 nm laser line and a EMCCD camera (iXon Ultra, Andor). Epi-illumination was used to bring fluorophores to single molecule blinking, and TIRF-illumination with 150 nm penetration depth was used for acquisition using Leica Application Suite × using high power mode (region of interest 18×18 µm^2^). The Alexa Fluor 647 channel was acquired prior to CF568 channel, and each channel was imaged for 4 minutes with 11 ms exposure per frame. 3D localization was performed using a custom written macro of ImageJ plug-in ThunderSTORM [48], and lateral drift was corrected using cross correlation. 647 and 568 channels were aligned using SMLMTools.jl [49] using TetraSpeck as a reference.

### 5.4 SuperResNET Merging and Filtering

We used the SuperResNET network analysis tool [9] to preprocess the SMLM point cloud data for all the datasets used. The program iteratively merges blinks within a predefined threshold to correct for multiple-blinking fluorophores; we used a threshold of 10 nm for the simulated dataset, a threshold of 20 nm for the PC3,PC3-CAVIN1 dataset, and a threshold of 14 nm for both channels in the Cav1-cavin-1 dataset. This is followed by a reduction of background noise not associated with any molecules through a filtering stage by comparing the degree graph of the localizations to that of a random graph with similar statistics. The degree graph was constructed using a proximity distance of 80 nm for both real datasets to determine the neighbors, while 60 nm was used for the simulated dataset. A parameter *α* is used to determine the filter cutoff. Given *D*_*r*_ as the mean degree in the random graph, the localizations that contain a degree less than or equal to the cutoff *α* · *D*_*r*_ are removed. For the PC3-CAVIN1 dataset, this value was set to *α* = 4. In the Cav1-cavin-1 dataset, the Cav1 channel used *α* = 0.5, while the cavin-1 channel used *α* = 1.95. This was determined by choosing an *α* that removed 95% from the background of manually segmented cells.

### 5.5 siMILe Hyperparameters

For the PC3, PC3–CAVIN1 dataset, within each condition and for each experimental replicate we reserved 2 cells for the validation set and 2 cells for the test set, using the remaining cells for training. Hyperparameters (bag size = 500, *σ* = 1000, *C* = 1000, *minacc* = 0.85) were selected by optimizing performance on the validation set and then fixed for evaluation on the held-out test set.

For the simulated dataset and its ablation study (siMILe, MILES+AE, MILES+SYM-C, MILES), we employ nested 5-fold cross-validation: an inner 5-fold grid search over the hyperparameter grid in Table 2, and an outer 5-fold loop to estimate held-out performance.

**Table 2.** Grid of candidate values for inner-loop tuning on the simulated dataset.

| Hyperparameter | Candidate values |
| --- | --- |
| Bag size | {5, 25, 50, 75, 100} |
| $\sigma$ | {0.1, 0.5, 1, 10, …, $10^6$} |
| $C$ | {0.01, 0.1, 0.5, 1, …, $10^4$} |
| $minacc$ | {0.5, 0.55, …, 0.95} |

## 6 Acknowledgements

This research was enabled in part by support provided by Westgrid (Cedar) and the Digital Research Alliance of Canada (alliancecan.ca). Funding of this project was kindly provided by the Canadian Institutes of Health Research (CIHR: PJT-175112) and the Natural Sciences and Engineering Research Council of Canada (NSERC: RGPIN-2019-05179, RGPIN-2020-06752). We acknowledge the helpful discussions of the members of the Nabi and Hamarneh labs in improving this project.

## 7 Supplementary Material

Source code: siMILe-code.zip

## Supplementary Material

### Supervised Learning Baseline

We first examined whether a standard supervised classifier could identify discriminative structures when trained with condition-level labels propagated to all instances. We trained a linear SVM with L1 regularization using the 30 SuperResNET blob features from the simulated dataset, labeling each blob according to its parent condition (A or B). The regularization parameter *C* was selected via grid search over {0.01, 0.1, 1, 10, 100, 1000}. To avoid data leakage from correlated blobs within the same simulated image, we performed file-level splitting, holding out entire images rather than individual blobs for testing.

The supervised classifier achieved only 55.2% accuracy in distinguishing conditions, marginally above random chance (50%), reflecting that the majority of blobs (type *c*) are common to both conditions and provide no discriminative signal. To assess whether the classifier could nonetheless identify discriminative structures, we examined its high-confidence predictions at varying thresholds (Table S1). At a confidence threshold of 0.7, the classifier achieved 90.2% precision but only 45.6% recall in identifying truly discriminative blobs (types *a* and *b*). Increasing the threshold to 0.8 improved precision to 100% but reduced recall to just 4.4%.

**Table S1.** Supervised baseline performance at varying confidence thresholds. Precision and recall are computed for identifying discriminative blobs (types *a* or *b*) among high-confidence predictions.

| Confidence Threshold | Precision | Recall |
| --- | --- | --- |
| 0.5 | 0.097 | 0.961 |
| 0.6 | 0.346 | 0.851 |
| 0.7 | 0.902 | 0.456 |
| 0.8 | 1.000 | 0.044 |

These results reveal fundamental limitations of the supervised approach for structure discovery. Even at the threshold achieving reasonable precision (0.7), recall remains below 50%, meaning the majority of discriminative structures go unidentified. This is particularly problematic when multiple types of discriminative structures exist with varying prominence in feature space; the classifier may preferentially identify the most distinctive structures while missing subtler but biologically important ones.

For biological applications where the goal is to identify all condition-specific structures, including those that may be less prominent or previously unknown, comprehensive recall is essential. This motivates the use of multiple instance learning, which explicitly models the relationship between bag-level labels and instance-level predictions. Moreover, siMILe’s adversarial erasing mechanism specifically addresses the challenge of comprehensive discovery by iteratively identifying discriminative structures beyond the most prominent ones.

### Guide to Users

siMILe’s applicability to other datasets depends on two separable components: feature extraction, which is domain-specific, and (2) the core MIL algorithm, which is domain-agnostic. The base algorithm (MILES), that siMILe extends, has been successfully applied across domains including drug discovery, computer vision, and histopathology. More importantly, our key innovations—adversarial erasing and symmetric classification—address fundamental MIL challenges (unknown witness rates and multi-condition discovery) that exist across domains, not just for Caveolae or even SMLM.

siMILe can analyze any dataset where: (1) there are structures/objects from two or more experimental conditions; (2) one can extract numerical features describing these structures; and (3) the goal is to identify condition-specific differences.

Practically, using siMILe requires having access to cells/images from 2 or more different biological conditions or experimental setups (e.g. wild-type/control, genetic mutation, over/under-expression of a protein, pathological condition) that need to be contrasted. From each of these cells/images, the user of siMILe should collect different features that describe the cellular structures, e.g. area/volumes, count, sphericity, border irregularity, etc. This feature extraction can be done with ImageJ, deep learning embeddings, or any domain-appropriate method. In this work, the interactive GUI-based SuperResNET software was the SMLM-specific feature extractor we adopted, which has been used to analyze various cellular structures in previous works [1, 2, 3].

Once features are collected from both conditions, they are collected into CSV files, where each row contains the features as well as an entry for the class (condition) of the cell, e.g. condition 1 vs. condition 2. siMILe expects the setting of the following parameters:

- *σ* **(sigma)** — Controls similarity bandwidth in the MILES embedding. Larger *σ* values create broader similarity, making the algorithm more conservative in labeling instances as discriminative. If over-iteration occurs, consider increasing *σ* substantially.
- **min_acc threshold** — Determines the stopping criterion for adversarial erasing iterations. The default threshold may be too permissive for some datasets. Increasing this value will terminate iterations earlier, preventing over-labeling of instances.
- **C parameter** — Controls regularization strength in the L1-SVM. High *C* values can cause overfitting to noise, leading to spurious discriminative patterns. Reducing *C* increases regularization and creates smoother decision boundaries.
- **bag_size** — Determines the number of instances per bag during MILES training. Smaller bags can amplify noise effects. Increasing bag size provides better statistical averaging and reduces sensitivity to outliers.

siMILe takes the CSV file and the parameters settings and runs its optimization to identify which sub-cellular structures, which rows of the CSV, are discriminant to each class or indiscriminatory.

Detailed usage instructions are provided on the siMILe GitHub repository: https://github.com/NanoscopyAI/siMILe.

**Table S2.**
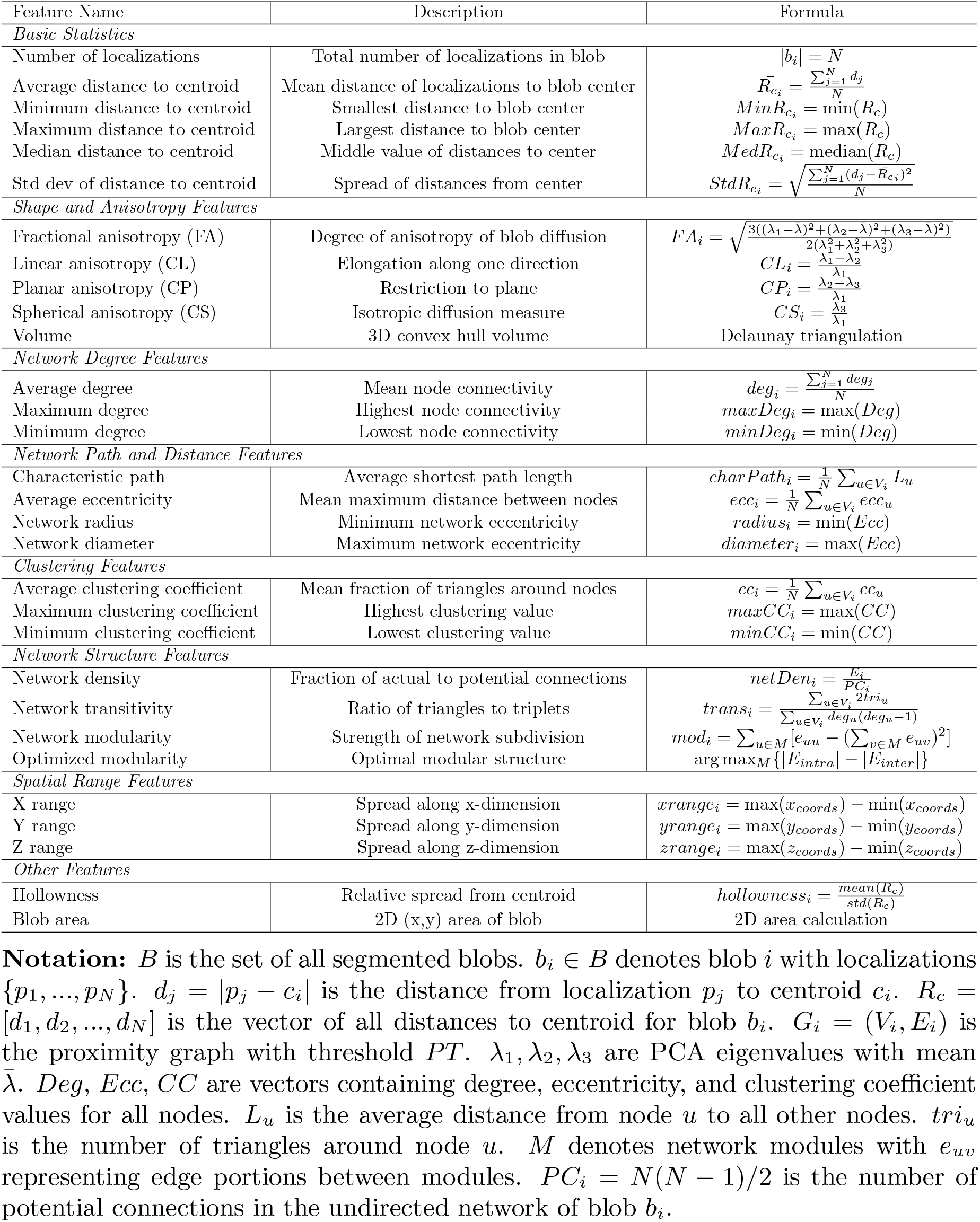
SuperResNET Blob Features.

### Algorithm Pseudocode

#### Algorithm S1 siMILe Pipeline

for identifying discriminative structures in SMLM data. Point clouds are segmented into blobs using SuperResNET, features are extracted for each blob, and the siMILe algorithm classifies each blob as discriminative to condition 0, discriminative to condition 1, or common to both.

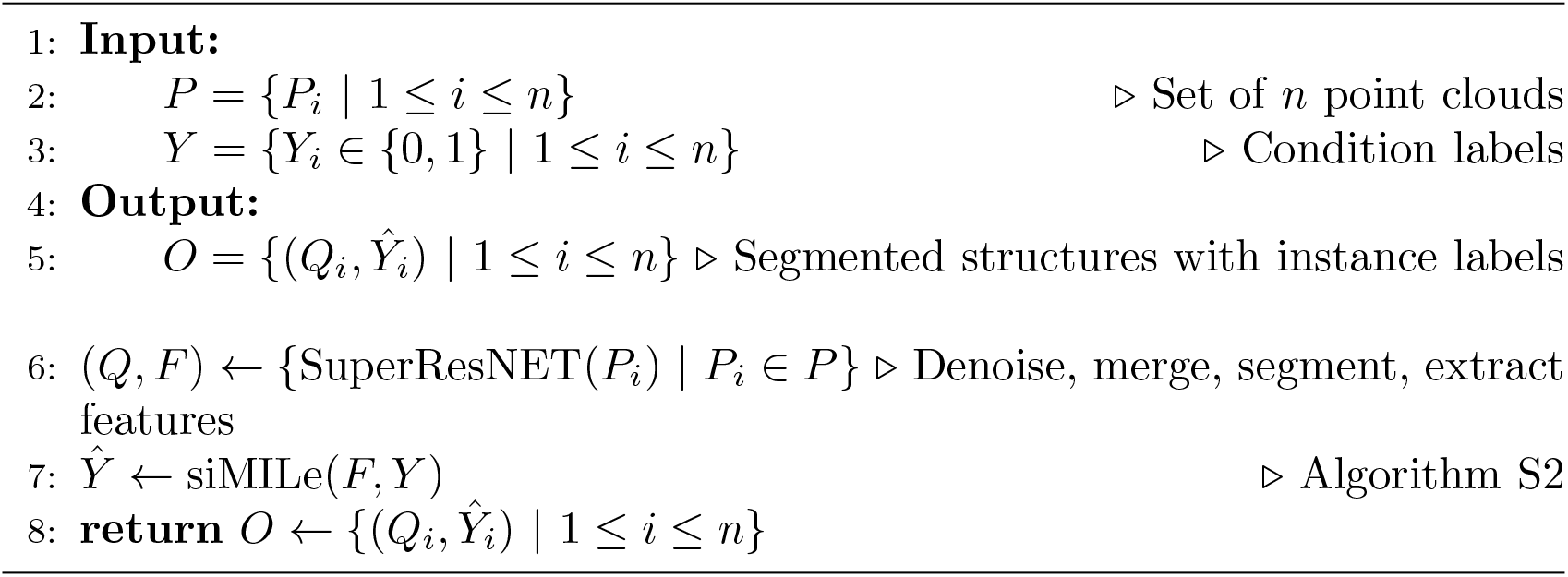

#### Algorithm S2 siMILe: Instance Classification with Adversarial Erasing

The algorithm iteratively trains a bag classifier, computes instance classification scores, and uses k-means clustering to identify discriminative instances from both conditions. Identified instances are removed (adversarial erasing) and the process repeats until bag classification accuracy falls below the threshold.

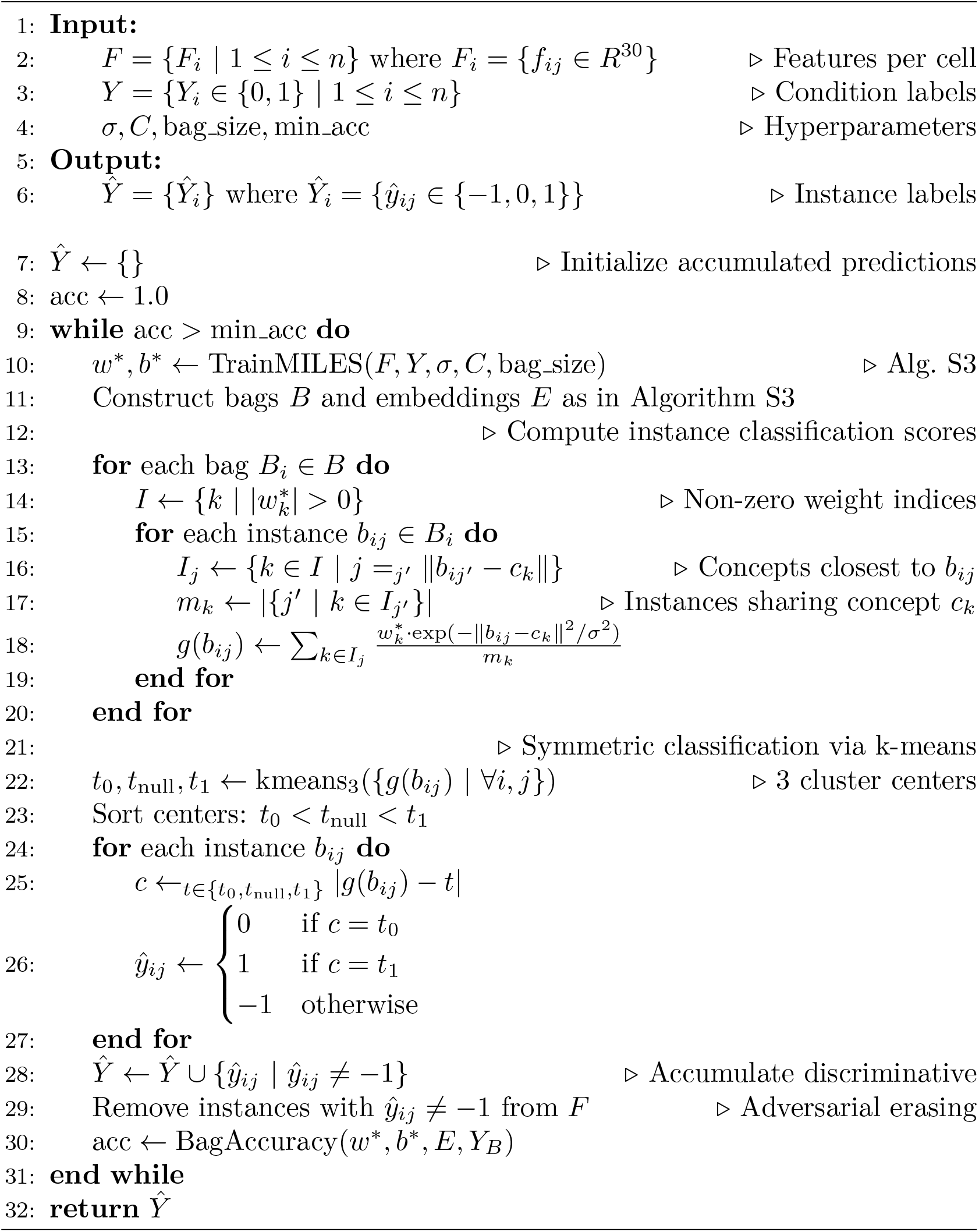

#### Algorithm S3 TrainMILES: Bag Classifier Training

Instances are pooled by condition and all serve as concepts. Bags are constructed by randomly grouping instances within each condition. Each bag is embedded by computing its maximum similarity to each concept, and an L1-regularized linear SVM is trained on these embeddings.

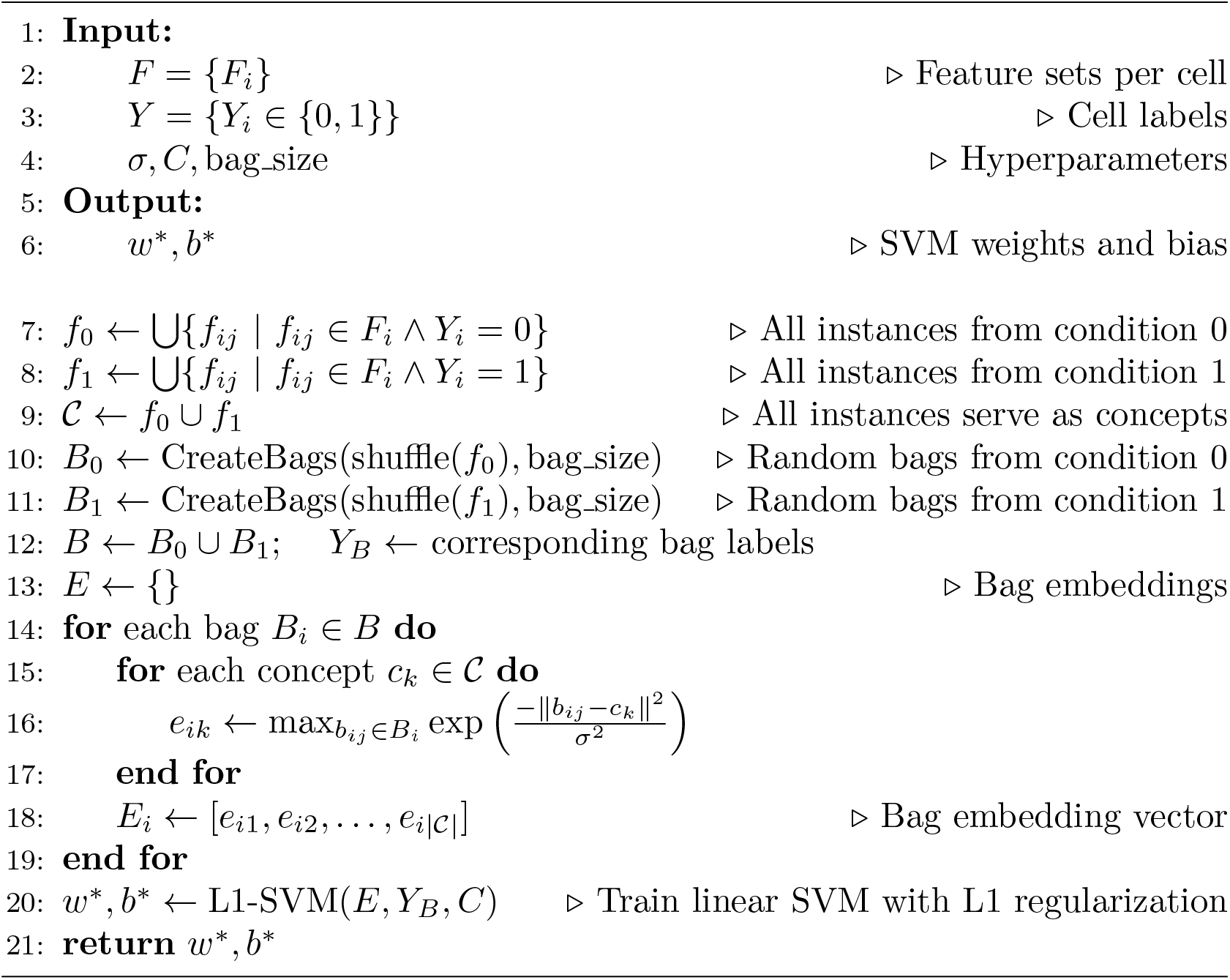

## References

[1] F. Balzarotti, Y. Eilers, K. C. Gwosch, A. H. Gynna, V. Westphal, F. D. Stefani, J. Elf, and S. W. Hell. Nanometer resolution imaging and tracking of fluorescent molecules with minimal photon fluxes. Science, 355(6325):606–612, 2017.

[2] Bo Huang, Wenqin Wang, Mark Bates, and Xiaowei Zhuang. Three-dimensional super-resolution imaging by stochastic optical reconstruction microscopy. Science, 319(5864):810–813, February 2008.

[3] Klaus C. Gwosch, Jasmin K. Pape, Francisco Balzarotti, Philipp Hoess, Jan Ellenberg, Jonas Ries, and Stefan W. Hell. Minflux nanoscopy delivers 3d multicolor nanometer resolution in cells. Nature Methods, 17(2):217–224, 2020.

[4] Nicholas Boyd, Eric Jonas, Hazen Babcock, and Benjamin Recht. Deeploco: Fast 3d localization microscopy using neural networks. bioRxiv, 2018.

[5] Leonhard Möckl, Anish R. Roy, Petar N. Petrov, and W. E. Moerner. Accurate and rapid background estimation in single-molecule localization microscopy using the deep neural network bgnet. Proceedings of the National Academy of Sciences, 117(1):60–67, 2020.

[6] I. M. Khater, I. R. Nabi, and G. Hamarneh. A review of super-resolution singlemolecule localization microscopy cluster analysis and quantification methods. Patterns (N Y), 1(3):100038, 2020.

[7] Yoonsuk Hyun and Doory Kim. Recent development of computational cluster analysis methods for single-molecule localization microscopy images. Computational and Structural Biotechnology Journal, 21:879–888, 2023.

[8] Yu-Le Wu, Aline Tschanz, Leonard Krupnik, and Jonas Ries. Quantitative data analysis in single-molecule localization microscopy. Trends in Cell Biology, 30(11):837–851, 2020.

[9] Ismail M. Khater, Fanrui Meng, Timothy H. Wong, Ivan Robert Nabi, and Ghassan Hamarneh. Super resolution network analysis defines the molecular architecture of caveolae and caveolin-1 scaffolds. Scientific Reports, 8(1):9009, Jun 2018.

[10] Patrick Lajoie, Jacky G Goetz, James W Dennis, and Ivan R Nabi. Lattices, rafts, and scaffolds: domain regulation of receptor signaling at the plasma membrane. J. Cell Biol., 185(3):381–385, May 2009.

[11] Ismail M. Khater, Qian Liu, Keng C. Chou, Ghassan Hamarneh, and Ivan Robert Nabi. Super-resolution modularity analysis shows polyhedral caveolin-1 oligomers combine to form scaffolds and caveolae. Scientific Reports, 9(1):9888, Jul 2019.

[12] Timothy H. Wong, Ismail M. Khater, Bharat Joshi, Mona Shahsavari, Ghassan Hamarneh, and Ivan R. Nabi. Single molecule network analysis identifies structural changes to caveolae and scaffolds due to mutation of the caveolin-1 scaffolding domain. Scientific Reports, 11(1):7810, Apr 2021.

[13] T H* Wong, I K* Khater, C Halgrimsson, YL Li, G* Hamarneh, and I R* Nabi. Molecular architecture of clathrin pit formation and inhibition defined by superresolution network analysis (superresnet). Submitted, 2024.

[14] Y L* Li, I M* Khater, C Hallgrimson, B Cardoen, TH Wong, G Hamarneh, and IR Nabi. Superresnet single molecule localization microscopy model-free network analysis achieves molecular resolution of nup96. Advanced Intelligent Systems, In press, 2024.

[15] Ibrahim M Khater, Stephanie T Aroca-Ouellette, Fan Meng, Ivan R Nabi, and Ghassan Hamarneh. Caveolae and scaffold detection from single molecule localization microscopy data using deep learning. PLoS ONE, 14(8):e0211659, 2019.

[16] Tamako Nishimura and Shiro Suetsugu. Super-resolution analysis of pacsin2 and ehd2 at caveolae. PLOS ONE, 17(7):1–13, 07 2022.

[17] Ivan R Nabi, Ben Cardoen, Ismail M Khater, Guang Gao, Timothy H Wong, and Ghassan Hamarneh. AI analysis of super-resolution microscopy: Biological discovery in the absence of ground truth. J. Cell Biol., 223(8), August 2024.

[18] Teun APM Huijben, Hamidreza Heydarian, Alexander Auer, Florian Schueder, Ralf Jungmann, Sjoerd Stallinga, and Bernd Rieger. Detecting structural heterogeneity in single-molecule localization microscopy data. Nature communications, 12(1):3791, 2021.

[19] Sobhan Haghparast, Sjoerd Stallinga, and Bernd Rieger. Detecting continuous structural heterogeneity in single-molecule localization microscopy data. Scientific Reports, 13(1):19800, 2023.

[20] Oren Z Kraus, Jimmy Lei Ba, and Brendan J Frey. Classifying and segmenting microscopy images with deep multiple instance learning. Bioinformatics, 32(12):i52–i59, 2016.

[21] M Sadegh Saberian, Kathleen P Moriarty, Andrea D Olmstead, Christian Hallgrimson, François Jean, Ivan R Nabi, Maxwell W Libbrecht, and Ghassan Hamarneh. Deemd: Drug efficacy estimation against sars-cov-2 based on cell morphology with deep multiple instance learning. IEEE Transactions on Medical Imaging, 41(11):3128– 3145, 2022.

[22] Francisco M. Castro-Macías, Pablo Morales-Álvarez, Yunan Wu, Rafael Molina, and Aggelos K. Katsaggelos. Sm: enhanced localization in multiple instance learning for medical imaging classification, 2024.

[23] Oded Maron and Tomás Lozano-Pérez. A framework for multiple-instance learning. Advances in neural information processing systems, 10, 1997.

[24] Yixin Chen, Jinbo Bi, and James Ze Wang. Miles: Multiple-instance learning via embedded instance selection. IEEE transactions on pattern analysis and machine intelligence, 28(12):1931–1947, 2006.

[25] Thomas G. Dietterich, Richard H. Lathrop, and Tomás Lozano-Pérez. Solving the multiple instance problem with axis-parallel rectangles. Artificial Intelligence, 89(1–2):31–71, January 1997.

[26] Stuart Andrews, Ioannis Tsochantaridis, and Thomas Hofmann. Support vector machines for multiple-instance learning. Advances in Neural Information Processing Systems, 15:561–568, 01 2002.

[27] Jun Wang and Jean-daniel Zucker. Solving the multiple-instance problem: A lazy learning approach. pages 1119–1126, 01 2000.

[28] Qi Zhang and Sally Goldman. Em-dd: An improved multiple-instance learning technique. In T. Dietterich, S. Becker, and Z. Ghahramani, editors, Advances in Neural Information Processing Systems, volume 14. MIT Press, 2001.

[29] Guoqing Liu, Jianxin Wu, and Zhi-Hua Zhou. pages Key instance detection in multi-instance learning. In Asian conference on machine learning, pages 253–268. PMLR, 2012.

[30] Oded Maron and Tomás Lozano-Pérez. A framework for multiple-instance learning. In M. Jordan, M. Kearns, and S. Solla, editors, Advances in Neural Information Processing Systems, volume 10. MIT Press, 1997.

[31] Corinna Cortes and Vladimir Vapnik. Support-vector networks. Machine learning, 20(3):273–297, 1995.

[32] Jeremy A Pike, Abdullah O Khan, Chiara Pallini, Steven G Thomas, Markus Mund, Jonas Ries, Natalie S Poulter, and Iain B Styles. Topological data analysis quantifies biological nano-structure from single molecule localization microscopy. Bioinformatics, 36(5):1614–1621, 2020.

[33] Michelle M. Hill, Michele Bastiani, Robert Luetterforst, Matthew Kirkham, Annika Kirkham, Susan J. Nixon, Piers Walser, Daniel Abankwa, Viola M.J. Oorschot, Sally Martin, John F. Hancock, and Robert G. Parton. Ptrf-cavin, a conserved cytoplasmic protein required for caveola formation and function. Cell, 132(1):113–124, January 2008.

[34] Lucas Pelkmans and Marino Zerial. Kinase-regulated quantal assemblies and kiss-and-run recycling of caveolae. Nature, 436(7047):128–133, July 2005.

[35] Timothy H. Wong, Ismail M. Khater, Christian Hallgrimson, Y. Lydia Li, Ghassan Hamarneh, and Ivan Robert Nabi. Superresnet – single-molecule network analysis detects changes to clathrin structure induced by small-molecule inhibitors (wong and khater: Joint first authors; hamarneh and nabi: Joint senior authors). Journal of Cell Science (JCS), 138(4):1–11, 2025.

[36] Sarah M. Smith and Corinne J. Smith. Capturing the mechanics of clathrin-mediated endocytosis. Current Opinion in Structural Biology, 75:102427, August 2022.

[37] Chao Zhang, Jialin Guo, Zixiao Liu, Xuhui Huang, Shiqi Dong, Chun Hu, and Junhai Xiao. Targeting clathrin-mediated endocytosis: recent advances in inhibitor development, mechanistic insights, and therapeutic prospects. RSC Medicinal Chemistry, 16(12):5843–5861, 2025.

[38] Defne Yarar, Clare M. Waterman-Storer, and Sandra L. Schmid. A dynamic actin cytoskeleton functions at multiple stages of clathrin-mediated endocytosis. Molecular Biology of the Cell, 16(2):964–975, February 2005.

[39] Lisa von Kleist, Wiebke Stahlschmidt, Haydar Bulut, Kira Gromova, Dmytro Puchkov, Mark J. Robertson, Kylie A. MacGregor, Nikolay Tomilin, Arndt Pechstein, Ngoc Chau, Megan Chircop, Jennette Sakoff, Jens Peter von Kries, Wolfram Saenger, Hans-Georg Kräusslich, Oleg Shupliakov, Phillip J. Robinson, Adam McCluskey, and Volker Haucke. Role of the clathrin terminal domain in regulating coated pit dynamics revealed by small molecule inhibition. Cell, 146(3):471–484, August 2011.

[40] Eric Macia, Marcelo Ehrlich, Ramiro Massol, Emmanuel Boucrot, Christian Brunner, and Tomas Kirchhausen. Dynasore, a cell-permeable inhibitor of dynamin. Developmental Cell, 10(6):839–850, June 2006.

[41] Tom Kirchhausen, Eric Macia, and Henry E. Pelish. Use of Dynasore, the Small Molecule Inhibitor of Dynamin, in the Regulation of Endocytosis, page 77–93. Elsevier, 2008.

[42] Bing Han, Alican Gulsevin, Sarah Connolly, Ting Wang, Brigitte Meyer, Jason Porta, Ajit Tiwari, Angie Deng, Louise Chang, Yelena Peskova, et al. Structural analysis of the p132l disease mutation in caveolin-1 reveals its role in the assembly of oligomeric complexes. Journal of Biological Chemistry, 299(4), 2023.

[43] Jason C Porta, Bing Han, Alican Gulsevin, Jeong Min Chung, Yelena Peskova, Sarah Connolly, Hassane S Mchaourab, Jens Meiler, Erkan Karakas, Anne K Kenworthy, et al. Molecular architecture of the human caveolin-1 complex. Science advances, 8(19):eabn7232, 2022.

[44] Arnold Hayer, Miriam Stoeber, Christin Bissig, and Ari Helenius. Biogenesis of Caveolae: stepwise assembly of large caveolin and Cavin complexes. Traffic, 11(3):361–382, 2010. Publisher: Wiley Online Library.

[45] Jeremy A. Pike, Abdullah O. Khan, Chiara Pallini, Steven G. Thomas, Markus Mund, Jonas Ries, Natalie S. Poulter, and Iain B. Styles. Topological data analysis quantifies biological nano-structure from single molecule localization microscopy. bioRxiv, 2018.

[46] Daniel J. Nieves, Jeremy A. Pike, Florian Levet, David J. Williamson, Mohammed Baragilly, Sandra Oloketuyi, Ario de Marco, Juliette Griffié, Daniel Sage, Edward A. K. Cohen, Jean-Baptiste Sibarita, Mike Heilemann, and Dylan M. Owen. A framework for evaluating the performance of smlm cluster analysis algorithms. Nature Methods, 20(2):259–267, February 2023.

[47] Logan R. Timmins, Milene Ortiz-Silva, Bharat Joshi, Y. Lydia Li, Fiona H. Dickson, Timothy H. Wong, Kurt R. Vandevoorde, and Ivan R. Nabi. Caveolin-1 promotes mitochondrial health and limits mitochondrial ros through rock/ampk regulation of basal mitophagic flux. The FASEB Journal, 38(1), December 2023.

[48] Martin Ovesný, Pavel Křížek, Josef Borkovec, Zdeňek Švindrych, and Guy M. Hagen. ThunderSTORM: a comprehensive ImageJ plug-in for PALM and STORM data analysis and super-resolution imaging. Bioinformatics, 30(16):2389–2390, 05 2014.

[49] Ben Cardoen. Smlmtools: A julia package for computational methods for single molecule localization / superresolution microscopy. February 2023.

## References

[1] Y L* Li, I M* Khater, C Hallgrimson, B Cardoen, TH Wong, G Hamarneh, and IR Nabi. Superresnet single molecule localization microscopy model-free network analysis achieves molecular resolution of nup96. Advanced Intelligent Systems, In press, 2024.

[2] Kailasam Mani, Nicolas Tardif, Olivier Rossier, Ismail M. Khater, Xuesi Zhou, Filipe Nunes Vicent, Radhakrishnan Av, Celine Gracia, Pamela Gonzalez Troncoso, Isabel Brito, Richard Ruez, Melissa Dewulf, Ghassan Hamarneh, Ivan Robert Nabi, Pierre Sens, Irina S Moreira, Gregory Giannone, Cedric M Blouin, and Christophe Lamaze. Remote control of cell signaling through caveolae mechanics. Technical Report biorxiv:2024.03.12.584716, Simon Fraser Universiry, 7 2025.

[3] Timothy H. Wong, Ismail M. Khater, Christian Hallgrimson, Y. Lydia Li, Ghassan Hamarneh, and Ivan Robert Nabi. Superresnet – single-molecule network analysis detects changes to clathrin structure induced by smallmolecule inhibitors (wong and khater: Joint first authors; hamarneh and nabi: Joint senior authors). Journal of Cell Science (JCS), 138(4):1–11, 2025.

